# The Genetics and Evolution of Eye Color in Domestic Pigeons (*Columba livia*)

**DOI:** 10.1101/2020.10.25.340760

**Authors:** Si Si, Xiao Xu, Yan Zhuang, Xiaodong Gao, Honghai Zhang, Zhengting Zou, Shu-Jin Luo

## Abstract

The avian eye color, generally referred to the color of the iris, results from both pigments and structural coloration. Avian iris colors exhibit striking interspecific and, in some domestic species, intraspecific variations, suggesting unique evolutionary and ecological histories. Here we tackled the genetic basis of the pearl (white) iris color in domestic pigeons (*Columba livia*), to elucidate the largely unknown genetic mechanism underlying the evolution of avian iris coloration. Using a genome-wide association study (GWAS) in 92 pigeons, we mapped the pearl iris trait to a 9 kb region and a facilitative glucose transporter gene *SLC2A11B*. A nonsense mutation W49X leading to a premature stop codon in SLC2A11B was identified as the causal variant. Transcriptome analysis suggested that SLC2A11B loss-of-function may downregulate the xanthophore-differentiation gene *CSF1R*, and a key gene *GCH1* involved in biosynthesis of pteridine, whose absence results in pearl iris. Coalescence and phylogenetic analyses indicated the mutation originated about 5,400 years ago coinciding with the onset of pigeon domestication, while positive selection was detected likely associated with artificial breeding. Within Aves, potentially impaired SLC2A11B was found in 10 species from six distinct lineages correlated to their signature brown or blue eyes. Analysis of vertebrate SLC2A11B orthologs revealed relaxed selection in the avian clade, consistent with the scenario that, during and after avian divergence from reptile ancestor, the SLC2A11B-involved development of dermal chromatophores likely degenerated due to feather coverage. Our findings provide new insight into the mechanism of avian iris color variations and the evolution of pigmentation in vertebrates.

## Introduction

Integumentary pigmentation of vertebrates plays an essential role in camouflage, sexual selection, communication, and thermoregulation (Hoekstra 2006; Hofreiter and Schoneberg 2010; Hubbard et al. 2010). The dynamic color change and diverse static integumentary pigmentation of poikilothermic vertebrates is mostly attributed to the neural crest-derived dermal chromatophores, which are generally divided into three main categories: xanthophore/erythrophore (yellow to red), iridophore/leucophore (irisescent color or white) and melanophore (black) (Grether et al. 2004; Kelsh 2004; Olsson et al. 2013).s

The eye color of a bird, usually referred to the color of the bird’s iris, derived from both pigmentation and structural coloration such as the diffraction of light. In birds, with the outer feather coverage masking the skin pigmentation, the dermal chromatophores may have undergone relaxed selection and was subject to evolutionary demise (Oliphant et al. 1992). However, the avian iris maintains the potential for complete development of all types of pigment cells that are comparable to the chromatophores in vertebrates, probably due to its external, exposed location where chromatophores are under constant selective pressure, thus remaining as a “pigment cell refugium” during avian evolution (Oliphant et al. 1992).

The pigment cells are located in the stroma, the anterior layer of iris consisted of loose vascular connective tissue. A bird’s eye color varies by the presence or absence of pigment cells in the iris and the content of pigments in the cells, and exhibits striking interspecific variations, ranging from black and dark brown, to brilliant colors, covering nearly the full spectrum of rainbow (Bond 1919; Oliphant 1987b; Oliphant et al. 1992). On the other hand, intraspecific color variation is rare and only common in domestic animals. Although the evolutionary drive shaping avian eye color remains largely unknown, recent studies had shed light on the possible coevolution of eye color and behavior or activity rhythm in birds (Craig and Hulley 2004; Davidson et al. 2014; Davidson et al. 2017; Passarotto et al. 2018). The iris color variation may reflect unique evolutionary history and ecological adaption, and thus provide a unique angle for understanding the avian radiation, as well as the evolution of pigmentation in vertebrates.

The domestic pigeon (*Columba livia)* was derived from its conspecific wild ancestor, the rock pigeon, and the origin of domestication is believed to occur around 5,000 years ago in the Mediterranean region (Darwin 1868; Price 2002; Driscoll et al. 2009). After the onset of domestication, pigeons have gone through intense selective breeding to have developed a wide array of phenotypic diversity within the species (Darwin 1868; Price 2002; Domyan and Shapiro 2017). Recent studies have revealed the molecular basis of some intraspecific variations in domestic pigeons such as plumage pigmentation, feather ornament, epidermal appendage, and navigation behavior, and highlighted the genes that might play important roles in avian evolution in these aspects (Shapiro et al. 2013; Domyan et al. 2014; Vickrey et al. 2015; Domyan et al. 2016; Gazda et al. 2018; Vickrey et al. 2018; Boer et al. 2019; Bruders et al. 2020; Shao et al. 2020).

The domestic pigeon exhibits three major types of iris color: the yellow to orange “gravel” eye (wild type, Fig. 1A), the white “pearl” eye (Fig. 1B), and the black “bull” eye (Bond 1919; Oliphant 1987b; Oliphant et al. 1992). The gravel and pearl irises in pigeons contain bright pigment cells with birefringent crystals in the anterior stroma tissue, while the bull eye results from a complete absence of stromal pigment cells (Oliphant 1987a). Crystalline guanine was identified in both gravel and pearl irises as the major pigment, and at least two yellow fluorescing pteridine materials were identified in the gravel iris leading to the yellow color tone (Oliphant 1987a). Therefore, the stromal pigment cell in the gravel iris was also referred as “reflecting xanthophore” (Oliphant 1987a). The irises of all newly hatched pigeons were bull-eyed dark, while those with gravel or pearl eyes gradually grew into brighter color after two to three months. Empirical breeding records indicated the pigeon pearl eye color is an autosomal recessive trait, denoted as the *Tr* locus, and the bull eye was associated with white feather (Hollander and Owen 1939). Similar stromal pigment cells and (or) developmental color changes were found in the irises of other avian species, suggesting the domestic pigeon is an ideal model for the evolution of avian pigmentation variations.

**Figure 1.**
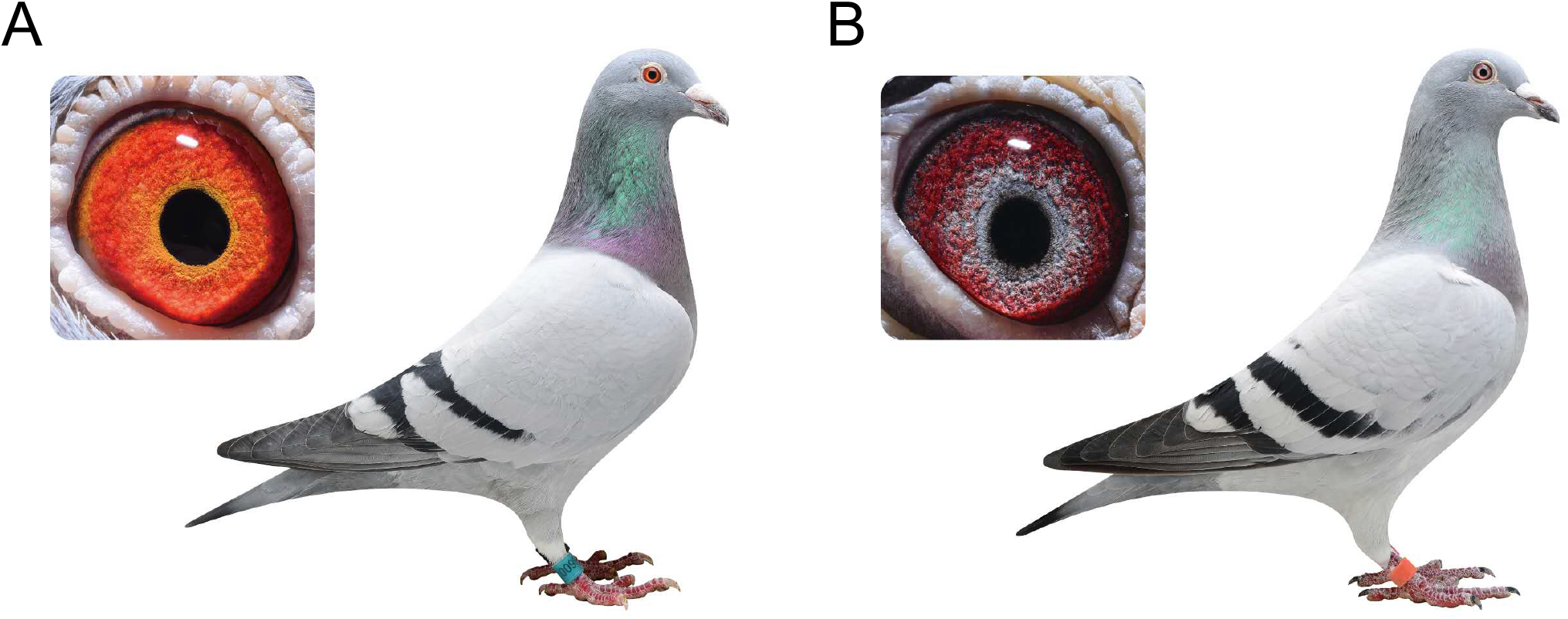
The iris color variation in domestic pigeons. (A) A pigeon with gravel eyes (wild type) exhibits bright orange iris color; (B) A pigeon with pearl eyes exhibits greyish white iris color. The red background in both gravel and pearl irises is from capillaries.

Here we investigated the genetic basis and origin of the bright irises (gravel and pearl eyes) in domestic pigeons. We identified the genetic mutation in SLC2A11B responsible for the iris color change from gravel to pearl, and found the pearl iris trait likely originated about 5,400 years ago as a domesticated trait under artificial selection. Evolutionary analyses elucidated a conserved role of SLC2A11B involved in avian eye color and associated its mutations with the lack of iris yellow pigmentation in various lineages of birds. Analysis of vertebrate SLC2A11B orthologs further revealed a relaxation of selection in the avian clade after its divergence from reptile ancestor, shedding new light on the origin and evolution of Aves coloration.

## Results

### *Tr* locus is mapped to a 9 Kb genomic region

A case-control genome-wide associate study (GWAS) based on whole genome resequencing was performed in domestic pigeons (racing homers) to identify the genomic region responsible for the bright iris phenotype. Whole genome sequencing data were retrieved from 49 gravel-eyed and 43 pearl-eyed individuals for an approximate 10× genome coverage each (Table S1). Of the 28,628,324 SNPs initially identified, 2,490,056 passed the genotype filtering and quality control and were used in subsequent analysis (Fig. S1). Significant signals of association (*P* < 2.01×10^−8^, after Bonferroni correction) were observed in 160 consecutive SNPs from the pigeon genome (Cliv_2.1, GenBank accession: GCA_000337935.2) scaffold AKCR02000030.1 (hereafter referred as scaffold 30), which spanned a single 341 Kb region associated with the pearl-eyed phenotype (minimum *P* = 1.07×10^−16^) (Fig. 2A, S2). No SNP from other scaffolds showed significant signal of association with the iris phenotypes (Fig. 2A, S2). In order to fine-map the *Tr* locus, haplotypes of the mapped region on scaffold 30 were built for each individual based on whole genome sequencing data. While wild-typed individuals exhibited multiple haplotypes in this region, a single 9 Kb haplotype (scaffold 30: 1894006-1903342) was shared by all 43 pearl-eyed pigeons, supporting it as a strong candidate for the *Tr* locus (Fig. 2B).

**Figure 2.**
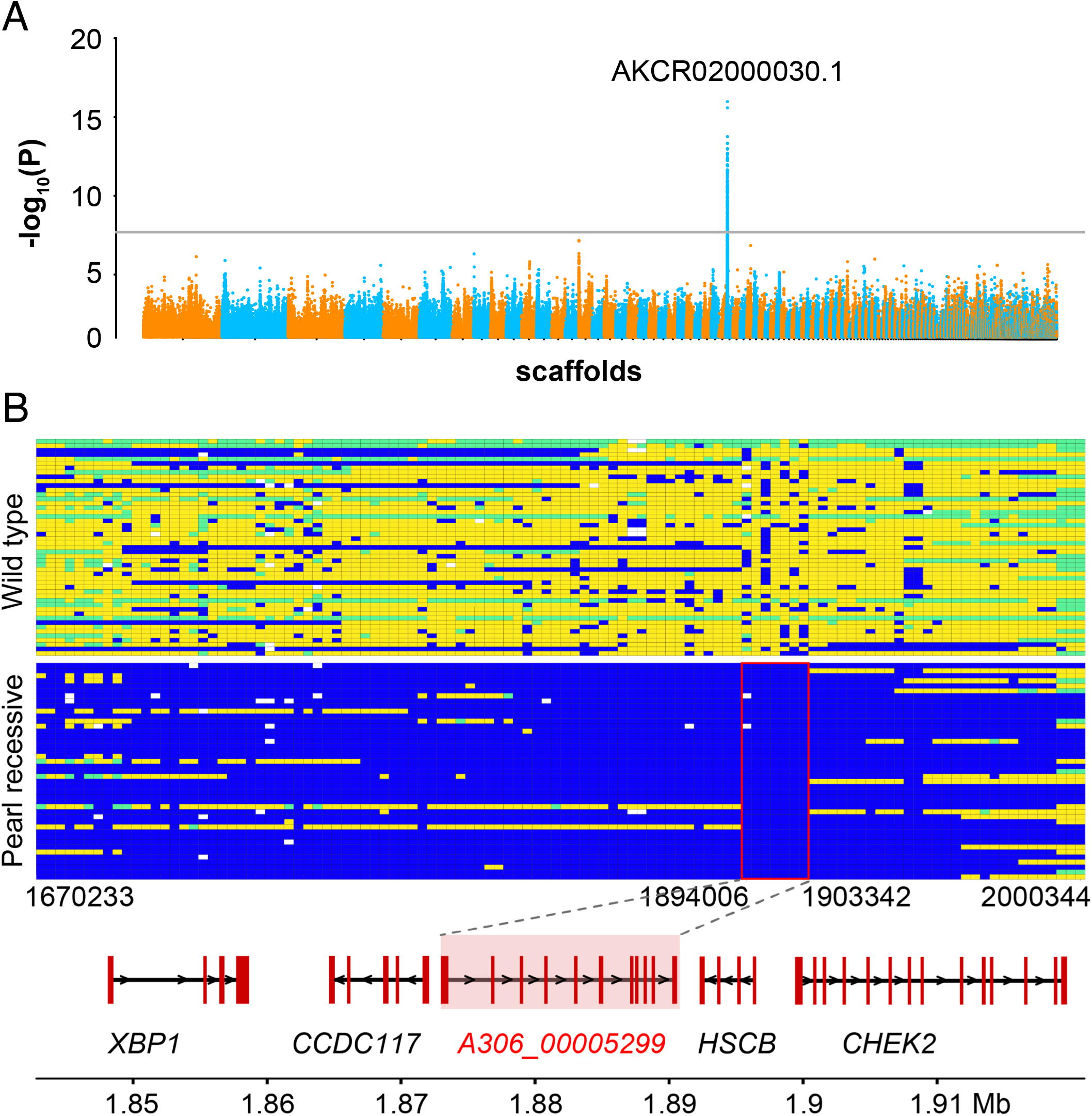
Genomic mapping of the *Tr* locus. (A) The Manhattan plot of the GWAS analysis for the *Tr* locus. The black line indicates the Bonferroni-corrected critical value (*P* = 2.01×10^−8^). The scaffold with the strongest association signal is marked. (B) The genotypes of 92 pigeons across the mapped region associated with the pearl eye. Each row represents one individual and each column represents an SNP/indel. Blue represents homozygosity for the major allele in pearl-eyed pigeons, yellow represents homozygosity for the minor allele in pearl-eyed pigeons, green represents heterozygosity, and white corresponds to missing data. A shared haplotype (scaffold 30: 1,894,006-1,903,342) was identified. Genes located in and around the shared region are specified.

### A premature stop codon of SLC2A11B causes the pigeon’s pearl eye color

The mapped 9 Kb interval of *Tr* locus contained only one protein-coding gene according to the gene annotation of the reference genome (Cliv_2.1): A306_00005299 (Fig. 2B). A306_00005299 was annotated as a 12-transmembrane transporter gene, solute carrier family 2 member 11-like (Fig. S3A-B), and was further identified as *SLC2A11B* (solute carrier family 2, facilitative glucose transporter, member 11b) based on gene homology and synteny in vertebrates (avian, reptile, and fish). The full length of pigeon *SLC2A11B* transcript was validated by RT-PCR from the iris tissue RNA of a gravel-eyed pigeon. *SLC2A11B* is known to play an essential role in the differentiation of leucophores and xanthophores in medaka fish (Kimura et al. 2014) and hence a strong candidate for the pearl eye color in domestic pigeons. In order to identify the causative mutation, screening of variations in the pigeon SLC2A11B coding region identified a G-to-A transition in the exon 3 that introduced a premature stop codon (W49X) and truncated the downstream 448 amino acids of SLC2A11B (Fig. 3A). Loss-of-function in SLC2A11B in medaka fish prevented the development of xanthophores and abolished the yellow pigment in leucophores at embryonic/larval stages (Kimura et al. 2014). SLC2A11B-knockout zebrafish also exhibited reduced yellow pigment due to the defect of the xanthophore differentiation (https://zmp.buschlab.org/gene/ENSDARG00000093395). The iris pigmentation in pearl-eyed pigeons resembles that of SLC2A11B loss-of-function in fish, supporting W49X of SLC2A11B as the causal mutation responsible for the pearl eye color. This nonsense mutation was further validated in 146 gravel-eyed and 146 pearl-eyed individuals. All gravel-eyed pigeons carried at least one wild-type allele, while 141 out of the 146 pearl-eyed individuals were homozygous for the mutant allele consistent with its recessive mode of inheritance. However, five pearl-eyed pigeons were heterozygous for the *SLC2A11B* p.W49X variant (Fig. 3B) and there was no other mutation found from the exons of *SLC2A11B*.

**Figure 3.**
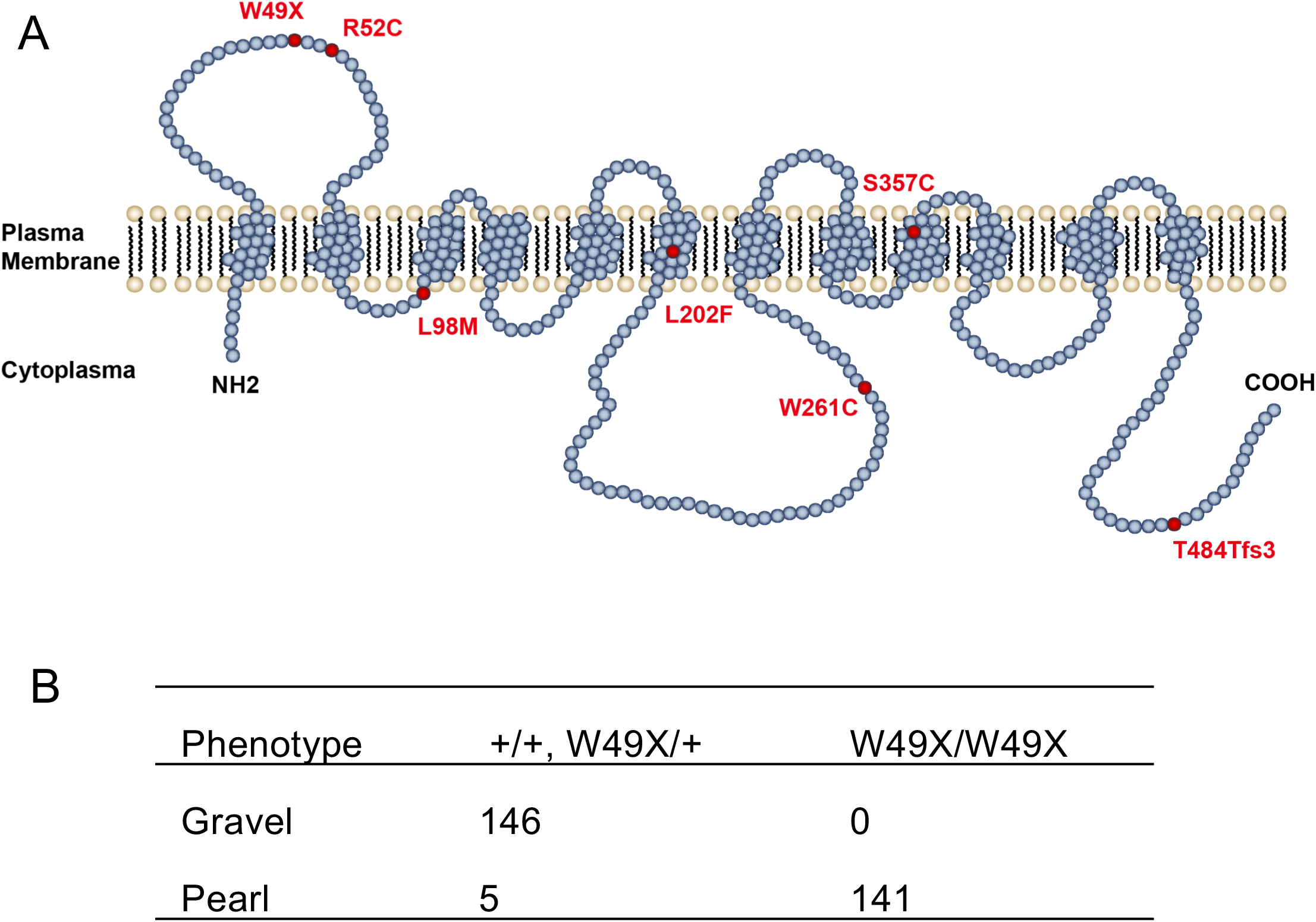
SLC2A11B mutations in avian species. (A) Schematic diagram for the 12 transmembrane SLC2A11B protein. The SLC2A11B mutations in birds were highlighted in red. (B) Validation of SLC2A11B W49X mutation in domestic pigeons.

### SLC2A11B W49X affects the stromal pigment cell differentiation in pigeons

The expression pattern of *SLC2A11B* was profiled in multiple tissues from nine wild-type domestic pigeons. Notably, the expression of *SLC2A11B* was high in iris tissue, but extremely low in skin and feather bud (Fig. 4A), indicating that the role of SLC2A11B in pigmentation may be restricted to the iris in pigeons. In addition, high expression of *SLC2A11B* was also observed in brain, retina, and muscle (Fig. 4A), suggesting its pleiotropic effect.

**Figure 4.**
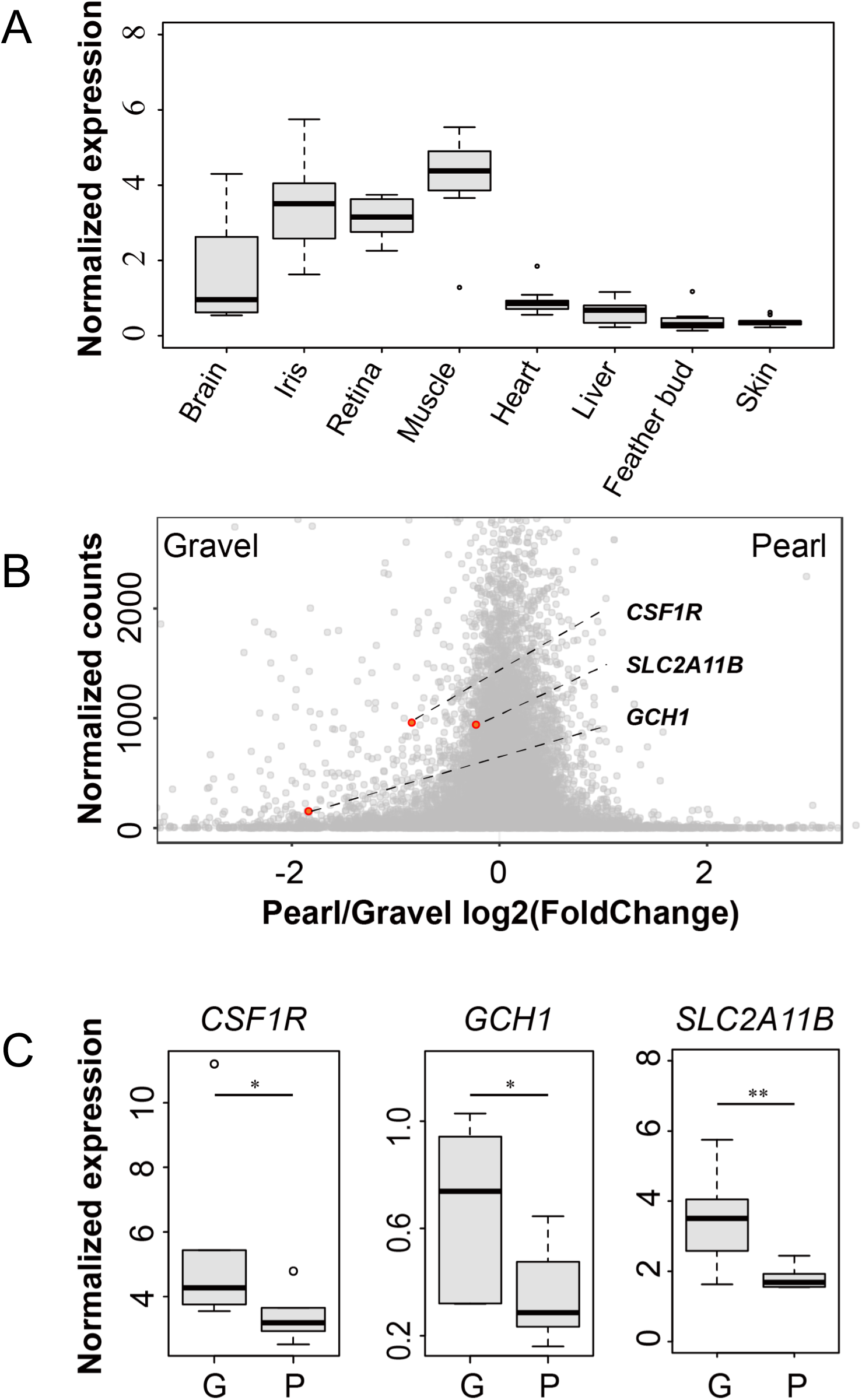
The expression profile of *SLC2A11B* in pigeon tissues and genes with differential expressions between pearl and gravel irises. (A) Relative expression of *SLC2A11B* in various tissues from wild-type pigeons with qPCR analysis. Boxes span the first to third quartiles, bars extend to the minimum and maximum observed values, black line indicates median, and circles represent the outlier values. (B) Normalized comparison of the *SLC2A11B* expression between the pearl and gravel irises by transcriptome sequencing. Differentially expressed genes involved in xanthophore-differentiation or pteridine synthesis are highlighted. (C) Validation of differentially expressed genes (*CSF1R, GCH1*, and *SLC2A11B*) between the pearl and gravel irises by qPCR analysis.

Transcriptome analysis of iris tissues from four gravel- and five pearl-eyed pigeons (Table S2) identified 337 differentially expressed genes (DEGs) (FDR < 0.05; Figs. 4B, S4, S5; Table S3). SLC2A11B was slightly downregulated by 1.2-fold in the pearl iris (Fig. 4B), likely caused by the nonsense-mutation-mediated RNA decay. Among these DEGs, we examined the genes involved in pteridine biosynthesize or xanthophore/leucophore differentiation. We identified that GTP cyclohydrolase 1 (*GCH1*), the gene encoding a rate-limiting enzyme in the pteridine pathway, was significantly downregulated (3.6 fold) in the iris tissues of pearl-eyed pigeons (Fig. 4B), consistent with the absence of yellow pigment. We also found colony-stimulating factor 1 receptor (*CSF1R*), a gene encoding a receptor tyrosine kinase required for the differentiation of the pteridine-containing xanthophore (Kelsh et al. 2009; Patterson et al. 2014; Singh and Nusslein-Volhard 2015), was significantly downregulated (1.8 fold) in pearl irises (Fig. 4B). Reduced expression of *SLC2A11B, GCH1*, and *CSF1R* were confirmed via quantitative RT-PCR (Fig. 4C), and jointly suggested that the loss-of-function variant of SLC2A11B may affect the differentiation of the xanthophore-like stromal pigment cells and pteridine biosynthesis machinery, resulting in the reduction of yellow pigment hence the pearl iris color in the pigeon.

### The pearl eye is a domesticated trait under artificial selection in pigeons

Genome data of fancy pigeons from representative breeds worldwide (Shapiro et al. 2013) was surveyed for the SLC2A11B mutation, to elucidate the evolution and origin of the pearl-eyed trait in pigeons (Table S4). The SLC2A11B W49X variant was found in 19 of the 36 breeds (Table S4). Such a widespread prevalence across pigeon breeds is consistent with a relatively ancient history of the pearl eye color in domestic pigeons.

To further investigate the origin of the SLC2A11B W49X mutation, a neighbor-joining gene phylogenetic tree was reconstructed based upon full-length *SLC2A11B* genic sequences from 140 domestic pigeons (36 fancy pigeons, 2 feral pigeons, and 102 racing homers) and one *C. rupestris*, (Fig. 5A). All haplotypes containing W49X mutation (*tr*^*mut*^) formed a monophyletic group nested within the wild-type (*Tr*^*+*^) clade (Figs. 5A, S6), indicating a derived state of the SLC2A11B variants originating from the wild type via one single mutation event. The derivative status of *tr*^*mut*^ in relation to *Tr*^*+*^ was also evident in the median-joining haplotype network (Fig. S6). In addition, the fact that all mutant haplotypes, while clustered together in the gene tree, spread across different breed sources and geographic regions (Fig. 5A) supports that the mutation is ancient and likely occurred prior to the establishment of modern pigeon breeds. We further estimated the time to the most recent common ancestor (TMRCA) for all *SLC2A11B* haplotypes bearing the W49X variant at about 5,400 years ago (95% CI:4,700-5800 years ago) (Fig. 5B), a time period consistent with the time of pigeon domestication estimated at more than 5,000 years ago.

**Figure 5.**
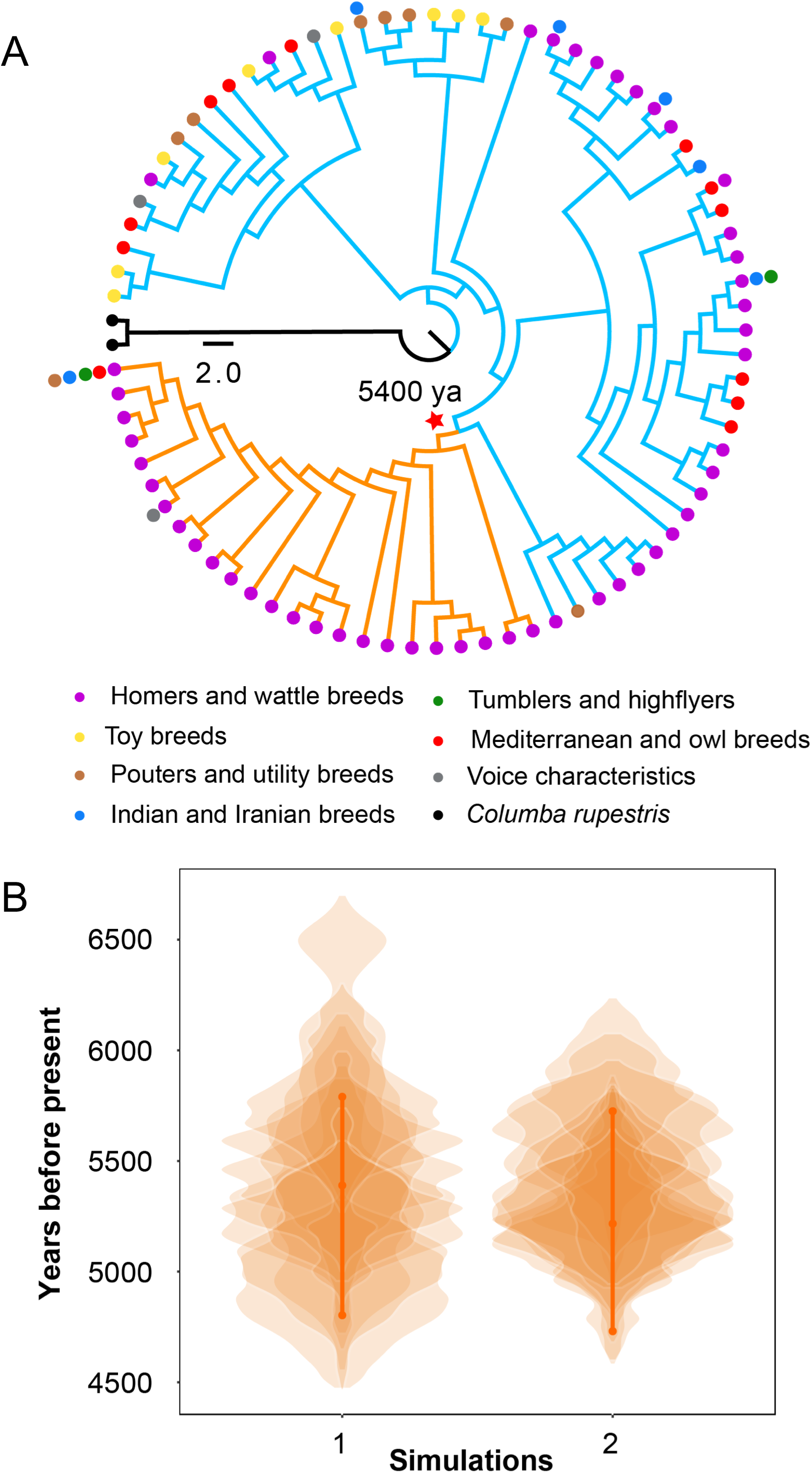
The origin of the pearl iris variant *tr*^mut^ in domestic pigeons. (A) Neighbor-joining (NJ) trees of 9 Kb nonrecombining *Tr* region based on 81 haplotypes from 141 unrelated individuals of 140 domestic pigeons (36 fancy pigeons, 2 feral pigeons, and 102 racing pigeons) and 1 *Columba rupestris* (outgroup). All nodes had more than 60% support from 1000 bootstrap replicates. Orange branches indicate *tr*^mut^ haplotypes, blue branches indicate *Tr*^*+*^ haplotypes, black branches indicate the outgroup. Different circle colors represent the seven traditional breeds of domestic pigeons and outgroup. Red star indicates the origin time of *Tr* mutation. (B) Violin plots of posterior distributions of the estimated TMRCA for *Tr* locus in domestic pigeons with the generation time of one year. Two independent simulations were performed using the estimated recombination rates of 10 Kb and 50 Kb windows, respectively. In every simulation, 10 replicate MCMCs are plotted with transparency.

Signals of selection in pigeons for the pearl eye trait were tested based on 140 genome data through evaluation of integrated haplotype score (iHS), nucleotide diversity(*π*), Tajima’s D, and extended haplotype homozygosity (EHH). We first applied iHS analysis on the scaffold 30 where *Tr* locus is located, and identified continuous signals of positive selection at the *Tr* locus and its adjacent region (scaffold 30: 1.4-2.0 Mb) (Figs. 6A, S7A, S7B). Decreased nucleotide diversity and negative Tajima’s D were evident across the SLC2A11B haplotypes with W49X mutant (Fig. 6C, 6D), consistent with a selective sweep scenario. The mutant haplotypes exhibited longer homozygosity than the wild-type ones in the EHH analysis (Figs. 6B, S7C, S7D), in support of the presence of selection for the variant. These evidences jointly illustrated that the prevalence of pearl iris trait in pigeons was likely a result of artificial selection during pigeon domestication.

**Figure 6.**
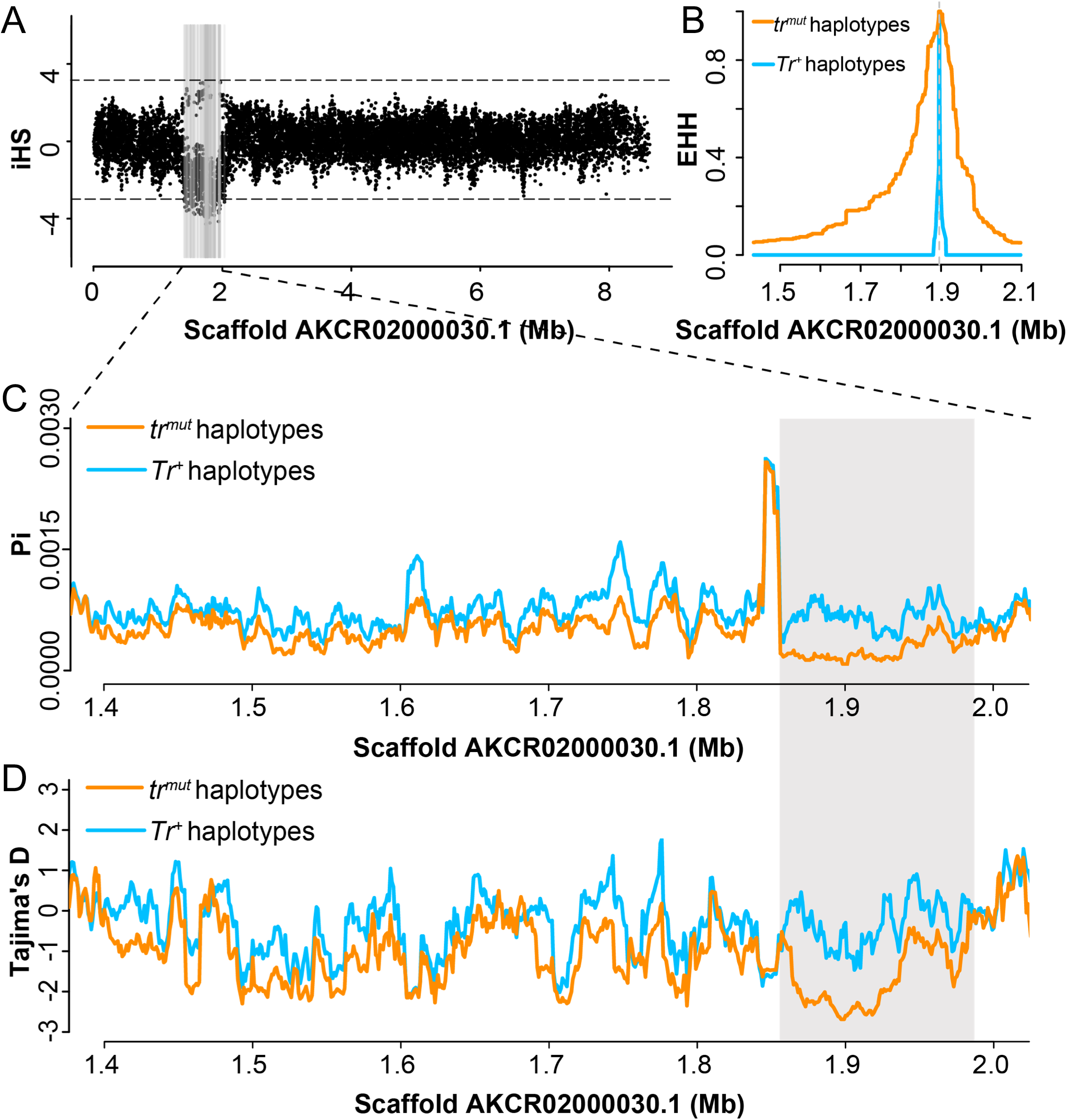
Selection of the pearl iris variant *tr*^*mut*^ in domestic pigeons. (A) The distribution of integrated haplotype score (iHS) values on scaffold AKCR02000030.1. The gray lines represent significance of the absolute iHS scores of 3 or greater. (B) The extended haplotype homozygosity (EHH) decay across the *Tr* locus region. Nucleotide diversity (*π*, Pi) (C) and Tajima’s D (D) were calculated in 10 Kb windows with 1 Kb step size for *tr*^*mut*^ and *Tr*^*+*^ haplotypes.

### SLC2A11B is involved in avian iris color variation

SLC2A11B plays an important role in the differentiation of xanthophore-like pigment cells in iris. Therefore, its defect may be responsible for the lack of pteridine and the iris color diversity in birds. Screening of sequence variations across *SLC2A11B* coding region in 70 avian species (Table S5) identified 230 nonsynonymous substitutions. After considering both evolutionary conservation and physicochemical constraint, five missense mutations that likely impaired the SLC2A11B function (with a deleterious SIFT score lesser than 0.05) were detected in species from five evolutionary diverging avian lineages: the Puerto Rican parrot (*<u>Amazona vittata</u>*, <u>R52C</u>), ostrich (*Struthio camelus*, L98M), Lawes’s parotia (*Parotia lawesii*, L202F), kea (*Nestor notabilis*, W261C), and cuckoo roller (*Leptosomus discolor*, S357C) (Table S6, Fig 3A, S8). All five species exhibit brown or blue iris (Table S5), with no hint of the xanthophore-like pigment cells or pteridine.

We found a reading frameshift mutation fixed in five cormorant species (family Phalacrocoracidae) from three different genera, or, *Nannopterum auratus, N. brasilianus, N. harrisi, Urile pelagicus*, and *Phalacrocorax carbo* (Fig. 3A, S8). The mutation was validated in one additional *P. carbo* specimen via PCR and sequencing. The T484Tfs3 truncated SLC2A11B C-terminus by 12 amino acids, most of which were evolutionarily conserved sites and hence the deletion might impair the protein’s function. All five cormorant species are featured by their bright blue irises without color hue of yellow pigment. Notably, a study in one of the species, *N. auratus*, has indicated that neither pteridine nor purine was present in the iris, and its signature blue eye was presumed to be a structural color of the melanocyte-like cells, generated by the membrane-bound granules consisted of electron-dense rods (Oliphant et al. 1992).

The association between various SLC2A11B loss-of-function variants and the absence of the xanthophore-like pigment cells across distinct avian lineages, suggested an SLC2A11B-involved, likely conserved, iris coloration pathway during avian evolution.

### SLC2A11B was under relaxed selection during or after the avian divergence from its reptile ancestor

With the development of a full feather coverage in birds and/or its feathered dinosaur ancestors, the functional constraints over the skin coloration may be gradually lifted, followed by the degeneration of dermal chromatophores, leaving only the epidermal melanocytes accounting for feather pigmentation. It is likely that genes specialized in the development of xanthophore or iridophore, or, *SLC2A11B* in this case, could have experienced relaxation in natural selection, during or after the avian divergence from its ancestral reptile lineage.

In order to test this hypothesis, we analyzed SLC2A11B orthologs from 41 species including five major vertebrate clades: Aves (birds), Crocodilia (crocodiles), Testudines (turtles), Squamata (scaled reptiles), and Teleostei (ray-finned fishes). RELAX analysis (Wertheim et al. 2015) was conducted, in which a statistic ***k*** is calculated denoting the strength of selection on a “Test” set of branches, normalized by that on a “Reference” branch set. We set three different schemes of Test/Reference branch designation, involving only the basal (“basal”) / all internal (“internal”) / all (“clade”) branches of each group (Fig. 7). With each scheme, relaxation of selection is tested for branch(es) of each species group against corresponding branches of the other four groups. We expect that birds show relaxation (*k* < 1) as Test branches in *SLC2A11B*, while no such significance is observed in *TYR*.

**Fig 7.**
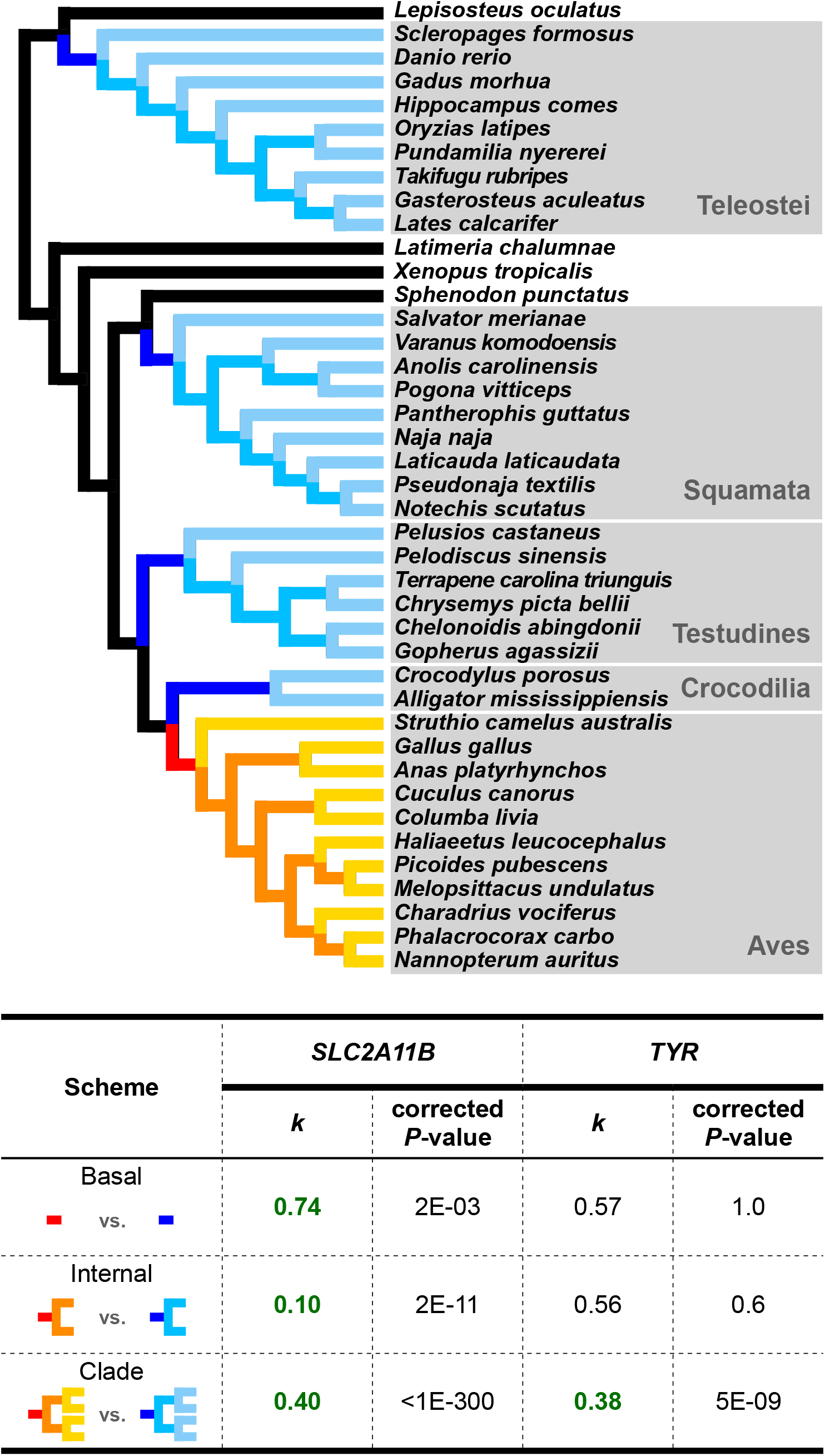
RELAX tests indicate consistent relaxation of selection in the Avian *SLC2A11B*, but not *TYR*. Three different schemes of Test/Reference branch designations were illustrated by color-labeled branches (red / dark blue for “basal”, red and orange / dark and intermediate blue for “internal”, all warm colors / all cool colors for “clade”) in the phylogeny and were tested separately (see Materials and Methods). The results of birds (warm-colored branches) as Test branches are shown in the table below (see Table S7 for more results). The ***k*** values significantly smaller than 1, which indicate relaxation of selection, are highlighted in green bold style. The *P*-values shown have been subject to Bonferroni multiple testing correction.

Relaxed selection in SLC2A11B was consistently detected along the avian clade (k < 1) under all three schemes of test/reference branch designations, but not in turtles or teleosts under any setting (Fig. 7, Table S7). Only weak signal of relaxation was partially detected in crocodilians under the basal scheme, and in squamates under the clade scheme, yet neither was consistent across all schemes (Table S7). On the other hand, no relaxed selection was consistently supported by all three schemes in *TYR*, the key melanogenesis gene that is expected to be evolutionarily conserved and under selective strength across all vertebrates (Fig. 7, Table S7). Overall, relaxation of selection in *SLC2A11B* in the avian clade, relative to other scenarios, is strongly supported, which is consistent with likely degeneration of *SLC2A11B*-involved development of dermal chromatophores in modern birds, due to the emergence of feather coverage during the evolutionary transition from the reptile ancestor.

## Discussion

Through whole genome sequencing and GWAS, we identified a nonsense mutation W49X of SLC2A11B responsible for the pigmentation change in the iris of pearl-eyed domestic pigeons. Most of the pearl-eyed individuals tested (141 out of 146) were homozygous for this mutation, consistent with its recessive mode of inheritance. One possible explanation for the exceptions of five pearl-eyed heterozygous pigeons might be additional variation(s) in SLC2A11B giving rise the same phenotype. Since no other mutation was identified from *SLC2A11B* coding regions, the additional mutation(s) involved in pearl iris color might locate in a different gene, or in the regulatory elements of *SLC2A11B*. Alternatively, the phenotype-genotype discordance in these five pearl-eyed individuals carrying one wild-type *SLC2A11B* allele might be due to mis-phenotyping. The iris of newly hatched domestic pigeons was black, and gradually exhibited bright colors after two to three months. Phenotypes of all the pigeons sampled in this study were recorded at about five months old when their iris color was usually stable. However, it is likely that the five heterozygous individuals might be younger than the others or there could be individual variance in the developmental stage of pigeons whose iris pigmentation change might be still in progress. Unfortunately, we are unable to trace back to these five discordant pigeons to confirm the potential phenotyping error, as they were lost during homing championship in the same year.

The W49X mutation would lead to truncation of about 90% amino acids of SLC2A11B, and was predicted to cause a total loss-of-function in the protein. In addition to iris, high expression of *SLC2A11B* was also evident in brain, muscle, and retina tissues, suggesting potentially multiple roles of the protein besides its involvement in pigmentation pathways. Therefore, it seems reasonable to expect consequences of the mutation other than iris pigmentation change. However, it seems the influence of *SLC2A11B* mutation in pigeons was restricted to the iris pigmentation, while no other abnormality was apparent in the pearl-eyed individuals. This is an observation consistent with the report that the loss-of-function of SLC2A11B in fishes only affected the pigmentation cells (Kimura et al. 2014). It is likely that the roles of SLC2A11B in non-pigmented cells of pearl-eyed pigeons (as well as other SLC2A11B mutants) could be compensated by the expression of other genes, mostly likely the other family members of solute carrier family 2.

Although the stromal pigment cell in the pigeon with gravel eyes is generally considered xanthophore-like, it shows a reflecting effect that is normally absent in a typical xanthophore. The stromal pigment cell in pigeons resembles the leucophore in medaka fish, in which the orange pigment is present at the larva stage, but diminishes in the SLC2A11B-defect mutant (Kimura et al. 2014). Therefore, it might be more appropriate to refer the pigeon iris stromal pigment cell as leucophore-like. Significantly reduced expression of *CSF1R*, a gene involved in xanthophore differentiation (Kelsh et al. 2009; Patterson et al. 2014; Singh and Nusslein-Volhard 2015), was detected in the pearl iris (*SLC2A11B* mutant), which indicated that *CSF1R* acts downstream of *SLC2A11B* in the regulatory network of xanthophore/leucophore differentiation. In addition, *GCH1*, the gene encoding the first and rate-limiting enzyme of pteridine biosynthesis (Ziegler 2003; Braasch et al. 2007), was also downregulated in the pearl iris. Since *GCH1* directly participates in the pteridine synthesis, it is plausible that *GCH1* is located in the far end of the regulatory pathway of xanthophore/leucophore differentiation, and triggers the pteridine production upon receiving the upstream signal.

While the precise timing of pigeon domestication is unclear, it is commonly accepted that the pigeon was first domesticated at least 5,000 years ago from the Fertile Crescent (Darwin 1868; Price 2002; Driscoll et al. 2009), and probably has served as a food resource for human since as early as 10,000 years ago (Darwin 1868; Driscoll et al. 2009; Shapiro and Domyan 2013). The estimated time of origin of *SLC2A11B* W49X mutation at about 5,400 years ago is in line with the commence of pigeon domestication, supporting that the pearl iris trait in domestic pigeons was a derived trait closely associated with the domestication process. Strong signals of positive selection were also detected, providing further evidence that the pearl iris trait has gone through artificial selection.

Even after thousands of years’ domestication, little morphological difference exists today between a wild-type domestic pigeon and its conspecific rock pigeon, it is thus reasonable to postulate that the pearl iris could serve as a domesticated marker for distinguishing a domestic individual from its wild counterpart during the early stage of domestication. In modern breeds, the reason for the selection of the pearl iris is more complicated. As the eye is one of the classic judging criteria for pigeons, the pearl iris has been preferred in many show breeds such as Show Homer, Runt, and Trumpeter. It is interesting that very high frequency of pearl iris is observed in performance breeds as well, such as Tumbler, Roller, and Highflier. Beside a random founder effect, breeders tend to hold an unfounded belief that the pearl eyes in a pigeon is somewhat associated with intelligence and hence better performance.

One landmark evolutionary transition setting the stage for the avian divergence from their dinosaur ancestors was the development of the feathery coverage that concealed their skin pigmentation. The dermal chromatophores would become functionally redundant and likely subject to evolutionary demise. In modern birds, only epidermal melanocyte retains throughout the body, playing an important role in feather pigmentation. The exception lies in the iris of the bird, which seems to maintain the full developmental potential for all types of chromatophores, and therefore has been proposed to be a pigment cell “refugium” (Oliphant et al. 1992).

In this context, our findings provide new insights into the mechanism underlying the evolutionary changes of the pigment cells in Aves. First of all, *SLC2A11B*, the gene first found involved in the differentiation of xanthophore and leucophore in fish, is absent in mammals but intact in birds. In pigeons, *SLC2A11B* exhibits a tissue-specific expression pattern, high in the iris and extremely low in the skin and feather bud. Evolutionary analysis of *SLC2A11B* orthologs in vertebrate supported that, relative to other reptiles and fishes, *SLC2A11B* was under a relaxation of selection in the avian clade. We proposed that during and after the avian divergence from its reptile ancestor, with the newly evolved feather coverage lifting selection pressure on underlying dermal coloration, the expression of *SLC2A11B*, as well as other specialized pigmentation genes, may gradually switch down or off in the dermis, thus resulting in the degeneration of the dermal chromatophores of the avian ancestor. In the iris, however, the expression of *SLC2A11B* and other pigmentation genes might still be functional and the pigment cells sustained.

The roles of iris pigmentation genes could have gone through a dynamic and complicated process during avian evolution. At the early stage of avian radiation, it is likely that the iris color of the ancient birds only carried limited diversity, when the evolutionary constraint over the genes involved in the development of the non-melanophore chromatophores were just released. Subsequently, mutations accumulated in the dermal pigmentation genes and the iris color diversity in birds may emerge, some of which might be subject to adaptive significance and selection along different avian lineages. Further studies on *SLC2A11B*, as well as other pigmentation genes in birds, would promise to illuminate the adaptative, functional, and evolutionary process of the avian coloration.

## Materials and Methods

### Sample collection

Feather samples of domestic pigeons were collected at the Xiangguan Pigeon Racing Club, Panjin City, Liaoning province, China, in 2017. This pigeon racing club recruited over 7,000 newly hatched pigeons from hundreds of breeders in the spring, and raised the racers in a uniform manner for about half a year until the championship in the fall. All pigeons selected in the sample set had the same wild-type feather color (blue), so as to exclude the potential influence of other pigmentation genes on the pigeon’s iris color. No more than two pigeons from the same breeder were used to ensure unrelatedness among individuals. Pigeons were sampled at about five to six months old, when the iris color became stable. The iris color phenotype of each pigeon was examined visually during the sampling and photographed. Four to six feathers with follicles were plucked from each individual and a total of 292 pigeons (146 with gravel eyes and 146 pearl eyes) from 237 breeders were gathered for the study. In addition, tissues of brain, iris, retina, muscle, heart, liver, feather bud, and skin, were collected from 14 domestic pigeons (nine with gravel eyes and five pearl and). Tissues samples were immediately submerged in RNALater reagent (Qiagen, Germany), stored in RNase-free 5 ml Eppendorf tubes at 4 °C overnight, and then transferred to −80 °C for long-term storage.

### DNA extraction

Genomic DNA from feather samples was extracted using a DNeasy Blood and Tissue Kit (QIAGEN, Valencia, California, USA) following the manufacturer’s instructions. DNA quantity and quality were examined using agarose gel electrophoresis, a Nanodrop spectrophotometer (Thermo Fisher Scientific, USA), and a Qubit fluorometer.

### RNA extraction

RNA was extracted from the brain, iris, retina, muscle, heart, liver, skin, and feather bud of nine gravel-eyed pigeons, and the iris of five pearl-eyed pigeons. The tissues were carefully removed from RNALater and homogenized in TRIzol™ Reagent (Invitrogen, USA). RNA was isolated following the manufacturer’s instruction and stored at −80 °C for further use. RNA degradation and possible contamination were monitored on 1% agarose gels.

### Whole-genome resequencing, read mapping, and SNP calling

A total of 92 pigeon individuals (49 gravel and 43 pearl) from 88 breeders were selected for whole genome sequencing (Table S1). The whole-genome resequencing was conducted at Mega Genomics Corporation, Beijing. For each genomic DNA extract, multiplexing library preparation with a uniquely tagged 6-bp sequence index was performed following the standard Illumina library construction protocol (Illumina, San Diego, California, USA). The libraries with average insert size 250-300 bp were sequenced using an Illumina Novaseq sequencer, which generated 150 bp paired-end reads, reaching an average sequencing depth of 16-fold coverage and producing an average of 16 Gb of raw sequencing data per individual (Table S1).

The adaptor sequences at both ends of the reads and bases with phred quality <30 were trimmed with Cutadapt v1.16 (Martin 2011). The processed reads were subsequently aligned to the domestic pigeon reference genome (Cliv_2.1 pigeon genome assembly) with the Burrows-Wheeler Aligner v0.7.17 with the options and default parameters (Li and Durbin 2010).

Sequence Alignment/Map (SAM) format files were first imported to SAMtools v1.7 for binary format conversion (SAM to BAM) and sorted by coordinates using the default options and parameters (Li et al. 2009; Li 2011). We then marked and removed optical or PCR duplicate reads, QC failure reads, unmapped reads, supplementary alignment reads, and non-primary aligned reads using SAMtools v1.7, with only the unique mapped reads retained (Table S1) (Li et al. 2009; Li 2011).

SNP and small indels calling was performed by GATK v3.8.1.0 and GATK v4.0.2.1 according to the GATK best practices manual with the default parameters (McKenna et al. 2010). Variant calling was performed with hard filters using GATK v4.0.2.1 and BCFtools v1.3.1 based on these filterExpression in VariantFilteration algorithm: FisherStrand (FS) >0.3, StrandOddsRatio (SOR) >2.0, RMSMappingQuality (MQ) <50.0, ReadPosRankSumTest (ReadPosRankSum) < − 0.05 (McKenna et al. 2010; Li 2011).

### LD estimation

Linkage disequilibrium (LD) decay was measured by correlation coefficients (r^2^) with PopLDdecay v3.40 (http://github.com/BGI-shenzhen/PopLDdecay, accessed 21 Dec. 2018) with the following parameters: -MAF 0.1, -MaxDist 600, -Miss 0.6. The LD decay was plotted as pairwise LD versus pairwise distance between SNPs with a maximum distance of 50 kb using PopLDdecay (Zhang et al. 2019).

### Gene mapping by genome-wide association study (GWAS)

SNPs and indels were filtered with the overall quality score (QUAL) greater than 20, the minor allele frequency (maf) greater than 0.05, the maximum missing genotype rates per variant (geno) greater than 0.1, and the maximum missing genotype rates per sample (mind) greater than 0.1. The resulting 2,490,056 SNPs from 92 individuals were used in genome-wide association analysis (GWAS) with PLINK v1.9 (Purcell et al. 2007), including 49 wild-type individuals set as the control group and 43 pearl-eyed individuals as the case group. Chi-square test was applied for differences between the case and control allele frequency distributions and the level of significance cutoff was set at 2×10^−8^ after Bonferroni correction. The Manhattan plot and QQ plot were plotted using the qqman R package (Turner 2018).

### Identification of causative mutation

The genotypes of all SNPs with a MAF greater than 0.1 and P-value less than 2×10^−8^ in the genomic regions with significant signals of GWAS were examined. A region with continuous homozygous genotypes shared by pearl-iris pigeons was considered as a candidate region, and the genes within the region were considered candidate genes. SNPs and indels from the candidate region were screened for the putative mutations associated with the gravel/pearl iris. After excluding the SNPs and indels in non-coding region, SNPs and indels leading to amino acid changes were examined for the evolutionary constraints at each affected amino acid residue site. The non-synonymous substitutions at conserved sites among reptile and avian species, or indels causing reading frame shifting or affecting conserved amino acid residues, were considered as the putative mutations.

### Causal mutation validation

The putative mutation was validated in an extended collection of unrelated domestic pigeons with confirmed iris color phenotypes. The sample set consisted of 146 gravel- and 146 pearl-iris pigeons from China, including the 92 abovementioned individuals used in GWAS analysis. Full coding exons of *SLC2A11B* were further sequenced in five pearl-iris individuals. The primer sets to amplify *SLC2A11B* exons (Table S8) were designed on the basis of the domestic pigeon genome assembly (Cliv_2.1). PCR, subsequent Sanger sequencing, and sequence analysis were performed following previously described procedures (Xu et al. 2013).

### Transmembrane model prediction

A 2D transmembrane model of SLC2A11B was constructed according to the schematic representation of the GLUT family of proteins in previous studies (Bryant et al. 2002). The transmembrane regions and orientation of SLC2A11B were predicted by TMpred (https://embnet.vital-it.ch/software/TMPRED_form.html) and TMHMM Server v.2.0 (http://www.cbs.dtu.dk/services/TMHMM/) (Hofmann 1993; Sonnhammer et al. 1998; Krogh et al. 2001).

### Transcriptome sequencing

Transcriptome sequencing of RNA extracts from the iris tissues of four gravel- and five pearl-eyed pigeons was conducted in Novogene Corporation, Beijing, China. The quality and purity of RNA were evaluated using the NanoPhotometer® spectrophotometer (IMPLEN, CA, USA). RNA concentration was measured using Qubit® RNA Assay Kit in Qubit® 2.0 Fluorometer (Life Technologies, CA, USA). The integrity of RNA was assessed with an RNA Nano 6000 Assay Kit of the Agilent Bioanalyzer 2100 system (Agilent Technologies, CA, USA).

A total amount of 1.5 μg RNA per sample was used for RNA library preparations. Sequencing libraries were generated using NEBNext® UltraTM RNA Library Prep Kit for Illumina® (NEB, USA) following manufacturer’s recommendations, and index codes were added to attribute sequences to each sample. Library quality was assessed on the Agilent Bioanalyzer 2100system. The libraries were sequenced on an Illumina HiseqXten platform and 150 bp paired-end reads were generated, which produced around 10 Gb clean data per sample.

### RNA-seq bioinformatics analysis

RNA-seq data were processed in one batch including case (five pearl irises) and control (four gravel irises) individuals. The reads were mapped to pigeon reference genome assembly (Cliv_2.1) using HISAT v2.1.0 and counted against the predicted gene models using HTSeq-count (Anders et al. 2015; Kim et al. 2015). The total number of aligned reads was normalized by gene length and sequencing depth for an accurate estimation of the expression level. These normalized read counts (TPM and FPKM) were used to represent the expression level of each gene, and differentially expressed genes were determined by DESeq2 (Love et al. 2014). The genes were sorted according to their log2-transformed fold-change values in DESeq2, and a hierarchical clustering algorithm in pheatmap R package was applied to generate the expression profiles of differently expressed genes (DEGs) (Kolde 2015). Analyses of gene ontology and KEGG pathway enrichment for 337 DEGs were performed in DAVID v6.8 (Table S9) (Huang et al. 2009b, a).

### Quantitative real-time PCR (qPCR) validation

Quantitative PCR was performed to identify *SLC2A11B* expression profile in the tissues of brain, iris, retina, muscle, heart, liver, feather bud, and skin from nine gravel-eyed pigeons, and to validate the differential expression of *SLC2A11B, CSF1R*, and *GCH1* between the five pearl and four gravel iris tissues. The obtained RNA was reversed-transcribed to cDNA using High Capacity cDNA Reverse Transcription Kit (Thermo Fisher Scientific, USA) according to the manufacture’s protocol. cDNA was amplified using intron-spanning primers (Table S8) for each target by Quantitative Real-time PCR, PowerUp™ SYBR® Green Master Mix (Applied Biosystems, USA) and implemented in QuantStudio® 3 Real-Time PCR instrument (Applied Biosystems, USA). Three replicates from each sample were performed to determine the mean value. Actin beta (*Actb*) was chosen as an internal reference for gene normalization (Vickrey et al. 2018). Experimental data were manually analyzed in a normalized expression comparative Ct (−2^ΔΔCt^) model. Results were compared by a Wilcoxon Rank Sum Test and differences in expression levels were considered statistically significant if *P* < 0.05.

### Haplotype inference

We downloaded from NCBI the published whole genome sequencing data of 36 fancy pigeons, two feral pigeons, 10 racing pigeons, and one hill pigeon (*Columba rupestris*) (Table S4), and integrated them into our genome data of 92 pigeons, to fine map the peal iris causal genes and mutations. VCF files containing genotypes of scaffold AKCR02000030.1 for all the 141 individuals were phased using SHAPEIT v2, with the following parameters: --burn 10 --prune 10 --main 20 --states 200 -- window 0.1 --rho 0.001 --effective-size 20000 --thread 70 (O’Connell et al. 2014). The phased haplotypes of the *Tr* locus were divided into two clusters: the mutant haplogroup *tr*^*mut*^ containing the nonsense mutation (W49X) and the wild-type haplogroup *Tr*^*+*^. The phased file was converted to fasta format and then used for summary statistics and phylogenetic analysis.

### Genetic diversity and selection analysis

The haplotypes spanning the entire AKCR02000030.1 was applied for the integrated haplotype score (iHS) analysis, which evaluates the extent of an excess of homozygosity around the ancestral or derived allele (Voight et al. 2006). Nucleotide diversity around *Tr* locus (*π*, the average pairwise differences), and Tajima’s D (a measure of the skew in the site frequency spectrum) were calculated in 10 Kb window with 1 Kb step size (Tajima 1989), with different combinations of sequence type, population, and *Tr*^*+*^ or *tr*^*mut*^ haplotypes. The extended haplotype homozygosity (EHH) score was implemented to validate whether the partial selective sweep occurred at the *Tr*^*+*^ and *tr*^*mut*^ haplotypes. EHH measures the relationship between the frequency of an interest allele and the amount of LD surrounding it and provides the probability that two randomly chosen chromosomes out of a population are homozygous between the core haplotype and the increasingly distant SNP (Sabeti et al. 2002). Once a focal marker is given, the *tr*^*mut*^ mutation in this case, the LD decay from the core haplotype were measured using increasingly distant SNPs. The genetic diversity and Tajima’s D calculations were performed in TASSEL v5.0 (Bradbury et al. 2007), and significance of differences were tested using a Wilcoxon Rank Sum Test with continuity correction in R. EHH and iHS tests were implemented with the rehh package in R (Gautier and Vitalis 2012).

### Phylogenetic analysis

Phylogenetic trees were built based on *tr*^*mut*^ and *Tr*^*+*^ haplotypes across the *SLC2A11B* genic region. Individuals with GQ less than 5 in *Tr* mutation site were removed, retaining a sample set of 140 pigeons consisted of 102 racing pigeons, 35 fancy pigeons, two feral pigeons, and one hill pigeon (*Columba rupestris*). A 9-Kb non-recombinant region was selected after visual examination, and 24 *tr*^*mut*^ and 57 *Tr*^*+*^ haplotypes were identified from the dataset. A minimum evolution (ME) phylogenetic tree was constructed from the 81 haplotypes using Kimura 2-parameter model and neighbor-joining (NJ) approach as implemented in PAUP v4.0a (Wilgenbusch and Swofford 2003). The reliability of the nodes in the NJ tree was assessed by 1,000 bootstrap iterations. *C. rupestris* (NCBI: SAMN01057534) was selected as outgroup. Phylogenetic trees were illustrated with the FIGTREE v1.3.1 package and modified manually.

A median-joining haplotype network was built based on *SLC2A11B* haplotypes from 102 racing pigeons, 35 fancy pigeon, and two feral pigeons. Haplotypes were reformatted using DnaSP v6.12.03 (Rozas et al. 2017), and constructed into network using the network approach in PopART software (Leigh and Bryant 2015).

### TMRCA estimation of the *tr*^*mut*^ haplotypes

To trace the origin of the domestic pigeon’s pearl iris mutation, the most recent common ancestor (TMRCA) for all *tr*^*mut*^ alleles was estimated using startmrca (Smith et al. 2018). which leverages both the recombination rates and the accumulation of new mutations on the targeted allele’s ancestral haplotype. Relative to other Approximate Bayesian Computation methods, the approach is based on a hidden Markov model and the assumption that the focal allele is subject to positive selection. Individuals homozygous for the *tr*^*mut*^ allele were set as the reference panel. The selected and reference panels were set at 100 and 40 each time. To obtain selection-onset time, independent Monte Carlo Markov (MCMC) chains were run 10 times, each with 200,000 iterations and the first 6,000 iterations discarded as burn-in. The one with the highest posterior probability was considered as the TMRCA estimate. To obtain confidence interval, we took the 2.5 and 97.5 quantiles of each resulting distribution, and calculated the recombination rates of the scaffold AKCR02000030.1 using an extremely fast open-source software package (FastEPRR) with nonoverlapping 10k and 50k window size respectively (Gao et al. 2016). We used the mutation rate of 1.42×10^−9^ per site per generation (Shapiro et al. 2013) and the generation time of one year.

### Identification of *SLC2A11B* variation in Aves

We downloaded genome data of 70 avian species from NCBI (Table S5) and extracted *SLC2A11B* coding sequences with blastn v2.7.1 (Camacho et al. 2009). *SLC2A11B* orthologs from two Crocodilia species (*Crocodylus porosus, Alligator mississippiensis*), six Testudines species (*Pelodiscus sinensis, Chelonoidis abingdonii, Chrysemys picta bellii, Terrapene Carolina triunguis, Pelusios castaneus, Gopherus agassizii*), and nine Squamata species (*Anolis carolinensis, Varanus komodoensis, Pogona vitticeps, Salvator merianae, Naja naja, Pseudonaja textilis, Pantherophis guttatus, Laticauda laticaudata, Notechis scutatus*) were downloaded from Ensembl database (http://asia.ensembl.org/index.html, Release 101). These 87 coding sequences were aligned using MUSCLE (codon) in MEGA v10.1.8 (Kumar et al. 2018) for variation detection. The missense variations at the evolutionary conserved amino acid sites were used to predict their potential impact on protein function with Sorting Intolerant From Tolerant (SIFT) (Sim et al. 2012).

### Detection of relaxed selection in the avian *SLC2A11B*

To test for relaxed or intensified selection of *SLC2A11B*, we selected 41 orthologs of the zebrafish *SLC2A11B* and *TYR* gene in the Ensembl database (http://asia.ensembl.org/index.html, Release 101), and aligned the CDSs according to the corresponding amino acid sequence by the L-INS-i algorithm in MAFFT v7.471 (Katoh and Standley 2013). CDS sequences of two cormorant species are truncated so that only the 1,434 bp before the nonsense substitution are retained. To obtain the tree topology of 41 vertebrate species, the initial tree was generated by http://timetree.org/ and modified according to the species tree topology reported in various phylogenetic studies (Hedges 2012; Near et al. 2012; Prum et al. 2015; Reeder et al. 2015). *TYR*, the key melanogenesis gene that is expected to be evolutionarily conserved and under selection across vertebrates, is used as control in the analysis.

The RELAX test implemented in the HYPHY package (Wertheim et al. 2015) takes some branches in the tree as Test branches and some others as Reference branches (there can be unclassified branches), and infer a parameter ***k***, which is the selection strength of both positive and negative selection (i.e. deviation of omega from 1) on the Test branches, divided by that on the Reference branches. Hence, if ***k*** is significantly smaller than 1, it means relaxed selection on Test branches relative to Reference branches; if ***k*** is significantly larger than 1, it suggests intensified selection. The level of significance of ***k*** is tested by a likelihood ratio test (LRT). Three different schemes of Test/Reference branch designation (basal, internal and clade) were applied: (1) the basal scheme only assigns the basal branches of each monophyletic group as Test/Reference branches; (2) the internal scheme assigns all internal branches of a monophyletic group as Test/Reference branches; and (3) the clade scheme assigns all (internal and external) branches within a clade as Test/Reference branches.

There are five major monophyletic groups in the 41 taxa involved in this analysis: Aves (birds), Crocodilia (crocodiles), Testudines (turtles), Squamata (scaled reptiles), and Teleostei (ray-finned fishes). For each scheme, we conduct RELAX test separately for each group, with corresponding branches of each group as Test branches against Reference branches consisting of respective branches in the other four groups. We did identical tests for the CDS of *SLC2A11B* and the negative control, *TYR*.

### Data availability

The sequencing data for domestic pigeons accessions from this study have been deposited in the NCBI Sequence Read Archive (BioProject ID: PRJNA682513).

## Acknowledgments

We thank J. -O. Gao for coordinating the sampling logistics, H. Meng, X. Sun, H. Yu, Y. -C. Liu, Y. -T. Xing for collecting samples, C. Xie for helpful discussion, and K. -W. Jiang for providing avian samples for validation, and all the pigeon owners for donating samples. The project was supported by the National Key Research and Development Program of China (2017YFF0210303), the National Natural Science Foundation of China (NSFC 31970537, 32070598), and the Peking-Tsinghua Center for Life Sciences.

## Author contributions

X. X. and S. -J. L. conceived the idea for this study. X. X. and S. -J. L. designed the study. S. S. and X. X. conducted experiments. S. S., X. X., and Z. Z. performed analysis. Y. Z. assisted with sampling and technical issues. X. L. and H. Z. provided *P. carbo* sample. S. S., X. X., Z. Z., and S. -J. L wrote the manuscript.

## Declaration of interests

The authors declare no competing interests.

## Notes

### Competing Interest Statement

The authors have declared no competing interest.

## References

Anders S, Pyl PT, Huber W. 2015. HTSeq-a Python framework to work with high-throughput sequencing data. Bioinformatics 31:166–169.

Boer EF, Van Hollebeke HF, Park S, Infante CR, Menke DB, Shapiro MD. 2019. Pigeon foot feathering reveals conserved limb identity networks. Dev. Biol. 454:128–144.

Bond CJ. 1919. On certain factors concerned in the production of eye colour in birds. J. Genet. 9:69–81.

Braasch I, Schartl M, Volff JN. 2007. Evolution of pigment synthesis pathways by gene and genome duplication in fish. BMC Evol. Biol. 7.

Bradbury PJ, Zhang Z, Kroon DE, Casstevens TM, Ramdoss Y, Buckler ES. 2007. TASSEL: software for association mapping of complex traits in diverse samples. Bioinformatics 23:2633–2635.

Bruders R, Van Hollebeke H, Osborne EJ, Kronenberg Z, Maclary E, Yandell M, Shapiro MD. 2020. A copy number variant is associated with a spectrum of pigmentation patterns in the rock pigeon (Columba livia). PLoS Genet. 16.

Bryant NJ, Govers R, James DE. 2002. Regulated transport of the glucose transporter glut4. Nat. Rev. Mol. Cell Biol. 3:267–277.

Camacho C, Coulouris G, Avagyan V, Ma N, Papadopoulos J, Bealer K, Madden TL. 2009. BLAST plus: architecture and applications. BMC Bioinformatics 10.

Craig AJFK, Hulley PE. 2004. Iris colour in passerine birds: why be bright-eyed? S. Afr. J. Sci 100:584–588.

Darwin. 1868. The Variation of Animals and Plants under Domestication, vol. 1. In: London: John Murray.

Davidson GL, Clayton NS, Thornton A. 2014. Salient eyes deter conspecific nest intruders in wild jackdaws (Corvus monedula). Biol. Lett. 10.

Davidson GL, Thornton A, Clayton NS. 2017. Evolution of iris colour in relation to cavity nesting and parental care in passerine birds. Biol. Lett. 13.

Domyan ET, Guernsey MW, Kronenberg Z, Krishnan S, Boissy RE, Vickrey AI, Rodgers C, Cassidy P, Leachman SA, Fondon JW, 3rd, et al. 2014. Epistatic and combinatorial effects of pigmentary gene mutations in the domestic pigeon. Curr. Biol. 24:459–464.

Domyan ET, Kronenberg Z, Infante CR, Vickrey AI, Stringham SA, Bruders R, Guernsey MW, Park S, Payne J, Beckstead RB, et al. 2016. Molecular shifts in limb identity underlie development of feathered feet in two domestic avian species. Elife 5:e12115.

Domyan ET, Shapiro MD. 2017. Pigeonetics takes flight: Evolution, development, and genetics of intraspecific variation. Dev. Biol. 427:241–250.

Driscoll CA, Macdonald DW, O’Brien SJ. 2009. From wild animals to domestic pets, an evolutionary view of domestication. Proc. Natl. Acad. Sci. U. S. A 106:9971–9978.

Gao F, Ming C, Hu WJ, Li HP. 2016. New Software for the Fast Estimation of Population Recombination Rates (FastEPRR) in the Genomic Era. G3-Genes Genomes Genetics 6:1563–1571.

Gautier M, Vitalis R. 2012. rehh: an R package to detect footprints of selection in genome-wide SNP data from haplotype structure. Bioinformatics 28:1176–1177.

Gazda MA, Andrade P, Afonso S, Dilyte J, Archer JP, Lopes RJ, Faria R, Carneiro M. 2018. Signatures of Selection on Standing Genetic Variation Underlie Athletic and Navigational Performance in Racing Pigeons. Mol. Biol. Evol. 35:1176–1189.

Grether GF, Kolluru GR, Nersissian K. 2004. Individual colour patches as multicomponent signals. Biol. Rev. 79:583–610.

Hedges SB. 2012. Amniote phylogeny and the position of turtles. Bmc Biology 10.

Hoekstra HE. 2006. Genetics, development and evolution of adaptive pigmentation in vertebrates. Heredity 97:222–234.

Hofmann K. 1993. TMbase-A database of membrane spanning proteins segments. Biol. Chem. Hoppe-Seyler 374:166.

Hofreiter M, Schoneberg T. 2010. The genetic and evolutionary basis of colour variation in vertebrates. Cell. Mol. Life Sci. 67:2591–2603.

Hollander W, Owen R. 1939. Iris pigmentation in domestic pigeons. Genetica 21:408–419.

Huang DW, Sherman BT, Lempicki RA. 2009a. Bioinformatics enrichment tools: paths toward the comprehensive functional analysis of large gene lists. Nucleic Acids Res. 37:1–13.

Huang DW, Sherman BT, Lempicki RA. 2009b. Systematic and integrative analysis of large gene lists using DAVID bioinformatics resources. Nat. Protoc. 4:44–57.

Hubbard JK, Uy JA, Hauber ME, Hoekstra HE, Safran RJ. 2010. Vertebrate pigmentation: from underlying genes to adaptive function. Trends Genet. 26:231–239.

Katoh K, Standley DM. 2013. MAFFT Multiple Sequence Alignment Software Version 7: Improvements in Performance and Usability. Mol. Biol. Evol. 30:772–780.

Kelsh RN. 2004. Genetics and evolution of pigment patterns in fish. Pigment Cell Res 17:326–336.

Kelsh RN, Harris ML, Colanesi S, Erickson CA. 2009. Stripes and belly-spots --a review of pigment cell morphogenesis in vertebrates. Semin Cell Dev Biol 20:90–104.

Kim D, Langmead B, Salzberg SL. 2015. HISAT: a fast spliced aligner with low memory requirements. Nat. Methods 12:357–360.

Kimura T, Nagao Y, Hashimoto H, Yamamoto-Shiraishi Y, Yamamoto S, Yabe T, Takada S, Kinoshita M, Kuroiwa A, Naruse K. 2014. Leucophores are similar to xanthophores in their specification and differentiation processes in medaka. Proc. Natl. Acad. Sci. U. S. A 111:7343–7348.

Kolde R. 2015. pheatmap: Pretty Heatmaps. R package version 1.0. 8. In.

Krogh A, Larsson B, von Heijne G, Sonnhammer ELL. 2001. Predicting transmembrane protein topology with a hidden Markov model: Application to complete genomes. J. Mol. Biol. 305:567–580.

Kumar S, Stecher G, Li M, Knyaz C, Tamura K. 2018. MEGA X: Molecular Evolutionary Genetics Analysis across Computing Platforms. Mol. Biol. Evol. 35:1547–1549.

Leigh JW, Bryant D. 2015. POPART: full-feature software for haplotype network construction. Methods Ecol. Evol. 6:1110–1116.

Li H. 2011. A statistical framework for SNP calling, mutation discovery, association mapping and population genetical parameter estimation from sequencing data. Bioinformatics 27:2987–2993.

Li H, Durbin R. 2010. Fast and accurate long-read alignment with Burrows-Wheeler transform. Bioinformatics 26:589–595.

Li H, Handsaker B, Wysoker A, Fennell T, Ruan J, Homer N, Marth G, Abecasis G, Durbin R, Proc GPD. 2009. The Sequence Alignment/Map format and SAMtools. Bioinformatics 25:2078–2079.

Love MI, Huber W, Anders S. 2014. Moderated estimation of fold change and dispersion for RNA-seq data with DESeq2. Genome Biol. 15.

Martin M. 2011. Cutadapt removes adapter sequences from high-throughput sequencing reads. EMBnet. journal 17:10–12.

McKenna A, Hanna M, Banks E, Sivachenko A, Cibulskis K, Kernytsky A, Garimella K, Altshuler D, Gabriel S, Daly M, et al. 2010. The Genome Analysis Toolkit: A MapReduce framework for analyzing next-generation DNA sequencing data. Genome Res. 20:1297–1303.

Near TJ, Eytan RI, Dornburg A, Kuhn KL, Moore JA, Davis MP, Wainwright PC, Friedman M, Smith WL. 2012. Resolution of ray-finned fish phylogeny and timing of diversification. Proc. Natl. Acad. Sci. U. S. A 109:13698–13703.

O’Connell J, Gurdasani D, Delaneau O, Pirastu N, Ulivi S, Cocca M, Traglia M, Huang J, Huffman JE, Rudan I, et al. 2014. A General Approach for Haplotype Phasing across the Full Spectrum of Relatedness. PLoS Genet. 10.

Oliphant LW. 1987a. Observations on the Pigmentation of the Pigeon Iris. Pigm. Cell Res. 1:202–208.

Oliphant LW. 1987b. Pteridines and Purines as Major Pigments of the Avian Iris. Pigm. Cell Res. 1:129–131.

Oliphant LW, Hudon J, Bagnara JT. 1992. Pigment Cell Refugia in Homeotherms - the Unique Evolutionary Position of the Iris. Pigm. Cell Res. 5:367–371.

Olsson M, Stuart-Fox D, Ballen C. 2013. Genetics and evolution of colour patterns in reptiles. Semin. Cell Dev. Biol. 24:529–541.

Passarotto A, Parejo D, Cruz-Miralles A, Aviles JM. 2018. The evolution of iris colour in relation to nocturnality in owls. J. Avian Biol. 49.

Patterson LB, Bain EJ, Parichy DM. 2014. Pigment cell interactions and differential xanthophore recruitment underlying zebrafish stripe reiteration and Danio pattern evolution. Nat. Commun. 5.

Price TD. 2002. Domesticated birds as a model for the genetics of speciation by sexual selection. Genetica 116:311–327.

Prum RO, Berv JS, Dornburg A, Field DJ, Townsend JP, Lemmon EM, Lemmon AR. 2015. A comprehensive phylogeny of birds (Aves) using targeted next-generation DNA sequencing. Nature 526:569–U247.

Purcell S, Neale B, Todd-Brown K, Thomas L, Ferreira MAR, Bender D, Maller J, Sklar P, de Bakker PIW, Daly MJ, et al. 2007. PLINK: A tool set for whole-genome association and population-based linkage analyses. Am J Hum Genet 81:559–575.

Reeder TW, Townsend TM, Mulcahy DG, Noonan BP, Wood PL, Sites JW, Wiens JJ. 2015. Integrated Analyses Resolve Conflicts over Squamate Reptile Phylogeny and Reveal Unexpected Placements for Fossil Taxa. PLoS One 10.

Rozas J, Ferrer-Mata A, Sanchez-DelBarrio JC, Guirao-Rico S, Librado P, Ramos-Onsins SE, Sanchez-Gracia A. 2017. DnaSP 6: DNA Sequence Polymorphism Analysis of Large Data Sets. Mol. Biol. Evol. 34:3299–3302.

Sabeti PC, Reich DE, Higgins JM, Levine HZP, Richter DJ, Schaffner SF, Gabriel SB, Platko JV, Patterson NJ, McDonald GJ, et al. 2002. Detecting recent positive selection in the human genome from haplotype structure. Nature 419:832–837.

Shao Y, Tian HY, Zhang JJ, Kharrati-Koopaee H, Guo X, Zhuang XL, Li ML, Nanaie HA, Tafti ED, Shojaei B, et al. 2020. Genomic and Phenotypic Analyses Reveal Mechanisms Underlying Homing Ability in Pigeon. Mol. Biol. Evol. 37:134–148.

Shapiro MD, Domyan ET. 2013. Domestic pigeons. Curr. Biol. 23:R302–303.

Shapiro MD, Kronenberg Z, Li C, Domyan ET, Pan H, Campbell M, Tan H, Huff CD, Hu H, Vickrey AI, et al. 2013. Genomic diversity and evolution of the head crest in the rock pigeon. Science 339:1063–1067.

Sim NL, Kumar P, Hu J, Henikoff S, Schneider G, Ng PC. 2012. SIFT web server: predicting effects of amino acid substitutions on proteins. Nucleic Acids Res. 40:W452–W457.

Singh AP, Nusslein-Volhard C. 2015. Zebrafish Stripes as a Model for Vertebrate Colour Pattern Formation. Curr. Biol. 25:R81–R92.

Smith J, Coop G, Stephens M, Novembre J. 2018. Estimating Time to the Common Ancestor for a Beneficial Allele. Mol. Biol. Evol. 35:1003–1017.

Sonnhammer EL, von Heijne G, Krogh A. 1998. A hidden Markov model for predicting transmembrane helices in protein sequences. Proc. Int. Conf. Intell. Syst. Mol. Biol. 6:175–182.

Tajima F. 1989. Statistical method for testing the neutral mutation hypothesis by DNA polymorphism. Genetics 123:585–595.

Turner SD. 2018. qqman: an R package for visualizing GWAS results using QQ and manhattan plots. J. Open Source Softw. 3:731.

Vickrey AI, Bruders R, Kronenberg Z, Mackey E, Bohlender RJ, Maclary ET, Maynez R, Osborne EJ, Johnson KP, Huff CD, et al. 2018. Introgression of regulatory alleles and a missense coding mutation drive plumage pattern diversity in the rock pigeon. Elife 7.

Vickrey AI, Domyan ET, Horvath MP, Shapiro MD. 2015. Convergent Evolution of Head Crests in Two Domesticated Columbids Is Associated with Different Missense Mutations in EphB2. Mol. Biol. Evol. 32:2657–2664.

Voight BF, Kudaravalli S, Wen X, Pritchard JK. 2006. A map of recent positive selection in the human genome. PLoS Biol. 4:e72.

Wertheim JO, Murrell B, Smith MD, Pond SLK, Scheffler K. 2015. RELAX: Detecting Relaxed Selection in a Phylogenetic Framework. Mol. Biol. Evol. 32:820–832.

Wilgenbusch JC, Swofford D. 2003. Inferring evolutionary trees with PAUP. Curr. Protoc. Bioinformatics:6.4. 1-6.4. 28.

Xu X, Dong GX, Hu XS, Miao L, Zhang XL, Zhang DL, Yang HD, Zhang TY, Zou ZT, Zhang TT, et al. 2013. The Genetic Basis of White Tigers. Curr. Biol. 23:1031–1035.

Zhang C, Dong SS, Xu JY, He WM, Yang TL. 2019. PopLDdecay: a fast and effective tool for linkage disequilibrium decay analysis based on variant call format files. Bioinformatics 35:1786–1788.

Ziegler I. 2003. The pteridine pathway in zebrafish: Regulation and specification during the determination of neural crest cell-fate. Pigm. Cell Res. 16:172–182.

