## Supplemental figures for "The Genetics and Evolution of Eye Color in Domestic Pigeons (*Columba livia*)"

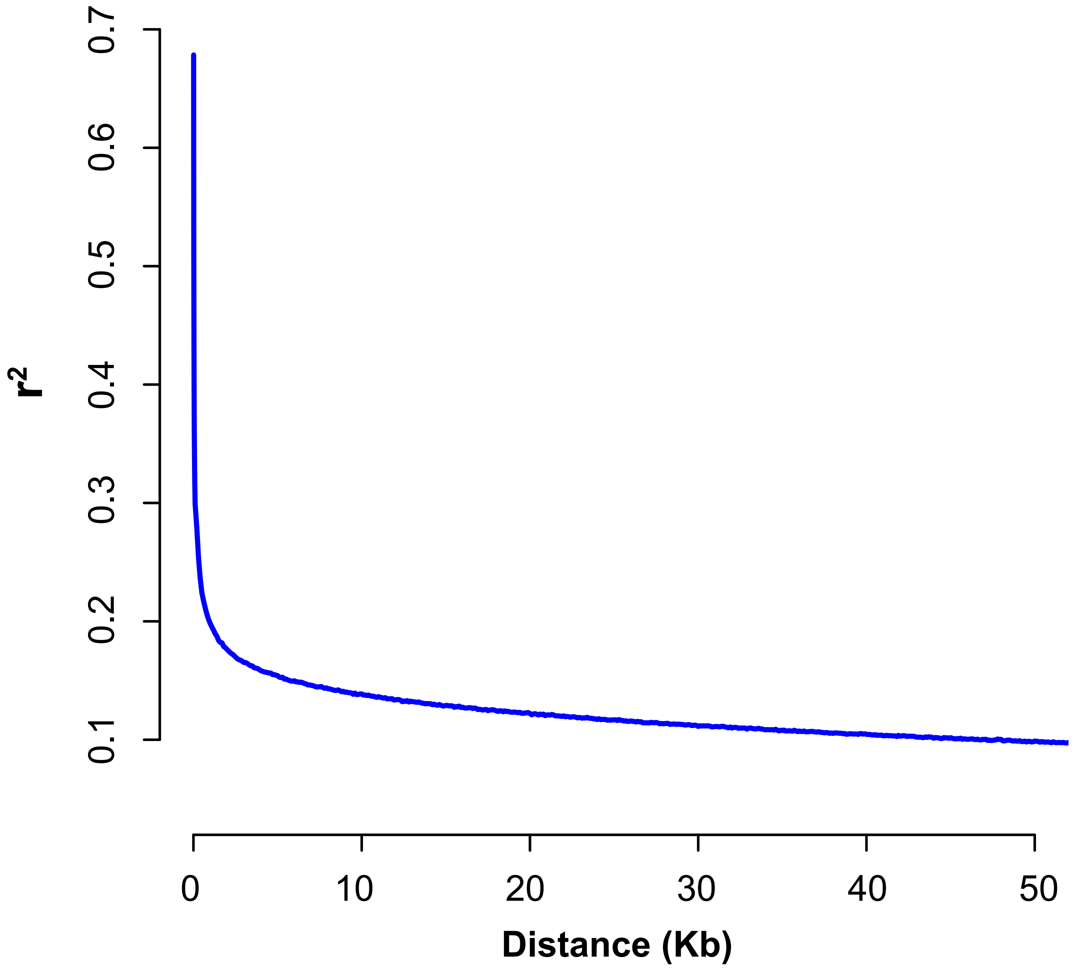


Figure S1. LD decay in racing pigeons. LD decay curve based on mean the genotype correlation coefficient ($r^2$) between common SNPs (minor allele frequency $\geq 0.1$). The threshold for "useful LD" with $r^2 < 0.2$ at distances beyond 0.9 Kb. Despite the rapid LD decay indicated that the clumps of SNPs were nearly or completely independent from each other, domestic pigeons were well suited for association-mapping methods considering on the sufficiently genome-wide marker density generated by SNPs calling.

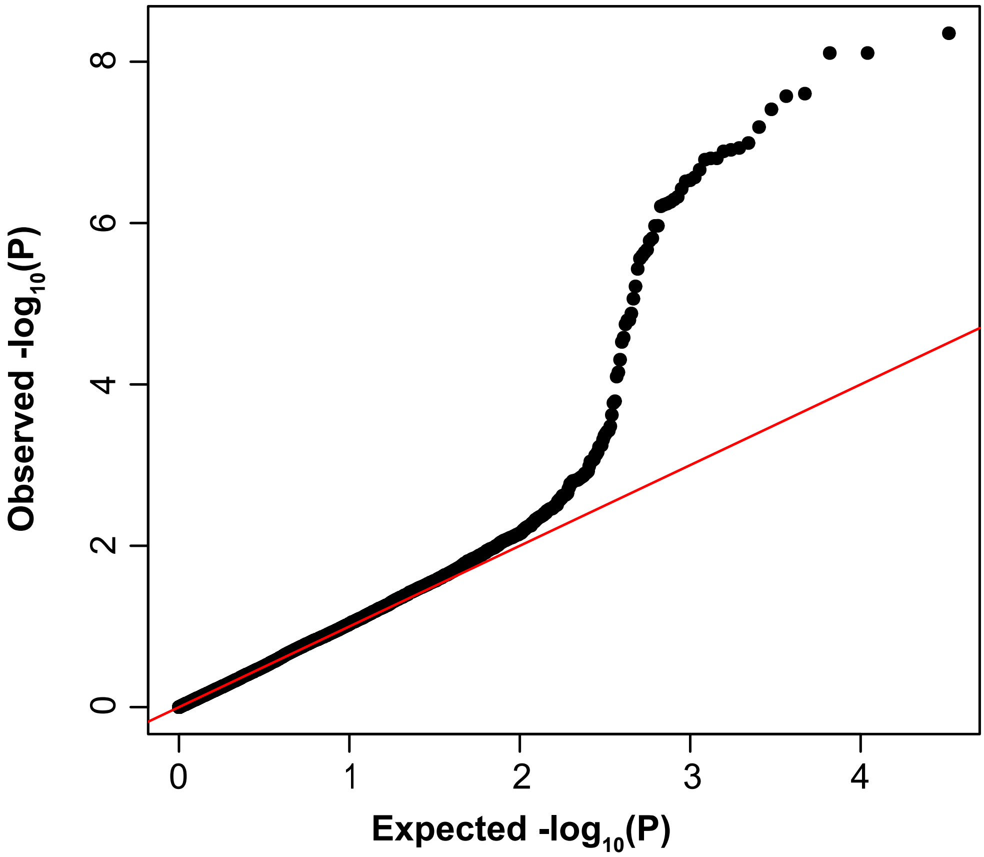


Figure S2. Quantile-Quantile (QQ) plot for GWAS. Observed versus expected quantiles of the genome-wide association p value shown in Figure 2A.

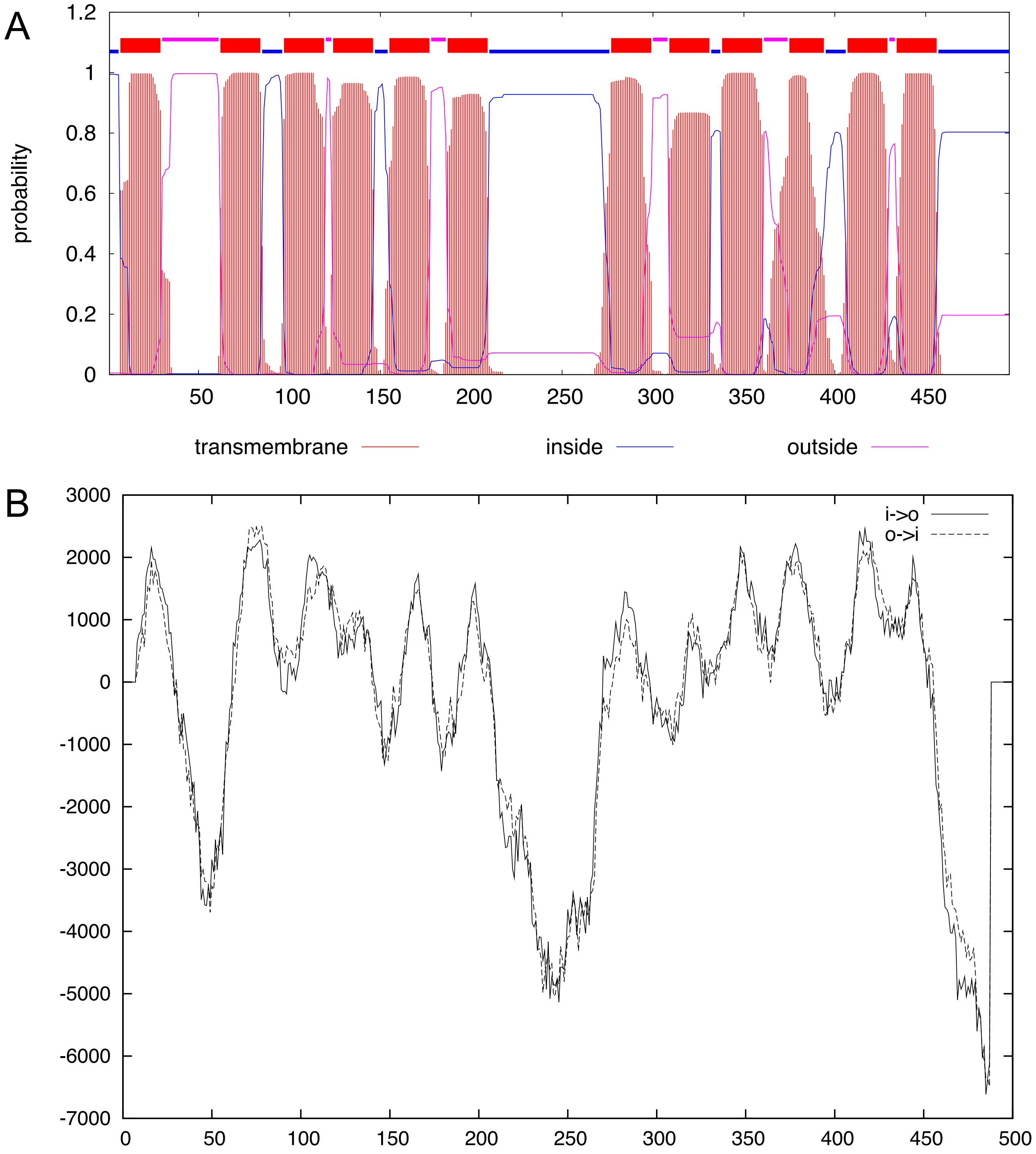


Figure S3. Prediction of transmembrane regions and orientation of SLC2A11B protein based on the whole SLC2A11B sequence of 496 amino acids. (A) TMHMM posterior probabilities of inside/outside/TM helix. The N-best prediction is displayed at the top where transmembrane regions are shown in red box. (B) Result output from TMpred server. The predicted transmembrane helices along with scores above 500 are considered significant. The solid and dashed line indicates inside to outside and outside to inside transmembrane helices respectively.

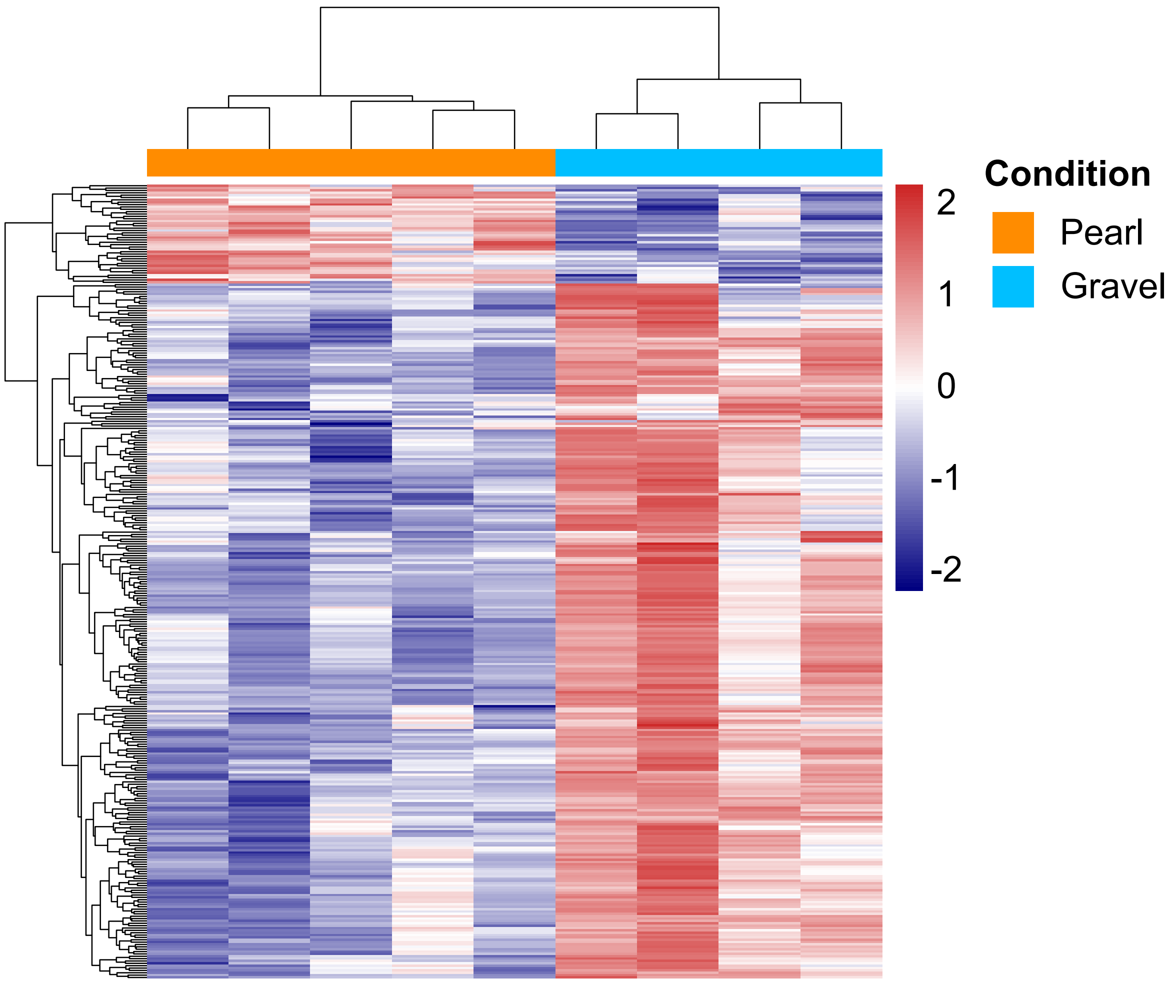


Figure S4. Gene expression profile between pearl and gravel irises by RNA-seq. The heatmap, hierarchically clustered into two group of genes, recapitulates a total of 337 differentially expressed genes (DEGs) between 5 pearl and 4 gravel irises, in which 295 and 42 genes were specifically upregulated in gravel and pearl irises, respectively. Columns are individual samples and rows indicate individual genes. Expression level is colored-coded from DESeq normalized counts, red represents high expression, blue represents lower expression.

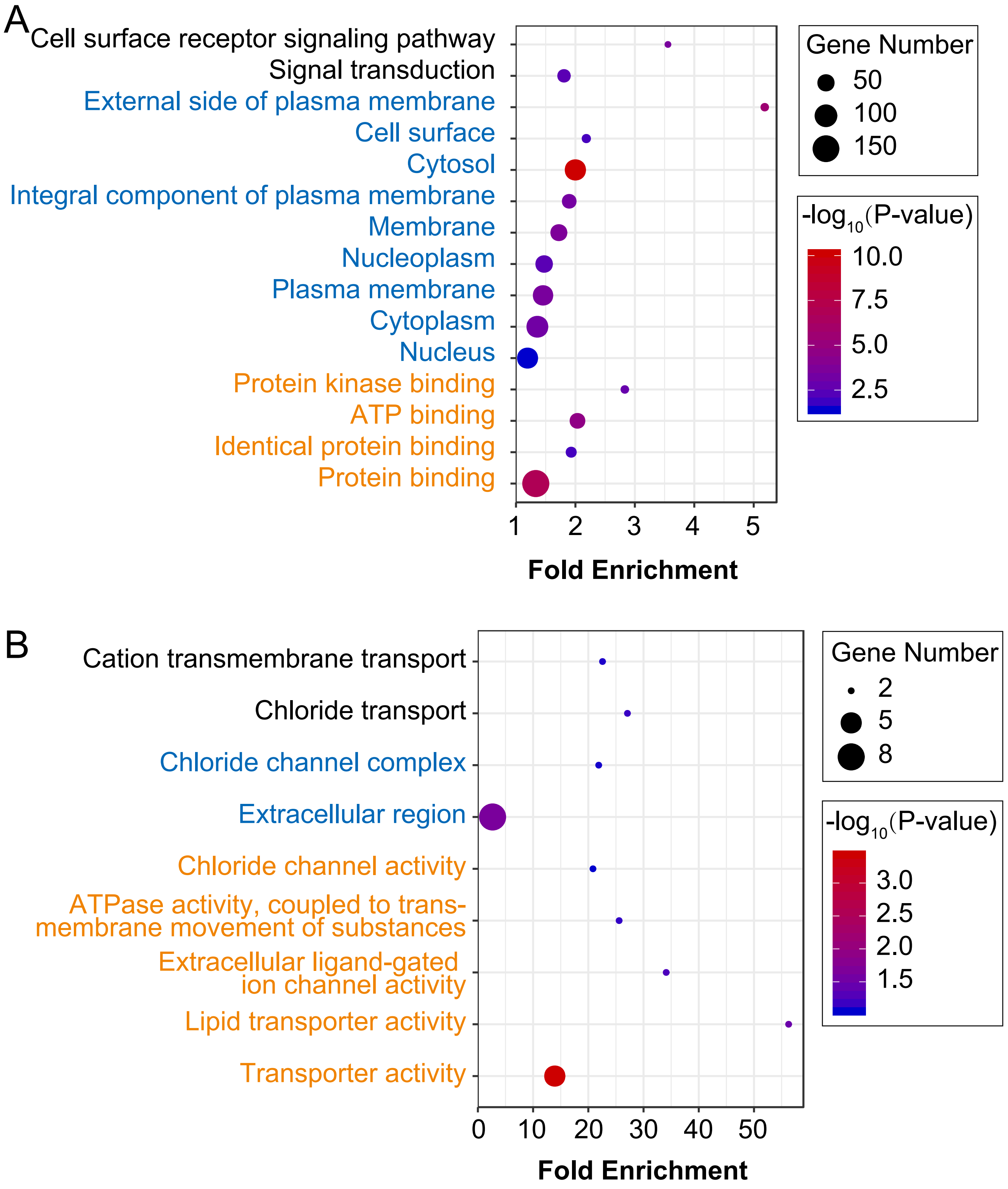


Figure S5. Functional analysis of differentially expressed genes (DEGs) based on RNA-seq data. (A) GO enrichment of 295 DEGs upregulated in gravel iris. (B) GO enrichment of 42 DEGs upregulated in pearl iris. The bubble diagrams show the degree of enrichment of Gene ontology (GO) terms in three categories. The orange, blue and black represent molecular function (MF), cellular component (CC), and biology process (BP) categories, respectively. Each bubble indicates a GO term, and the size of bubbles indicate the number of genes annotated to the GO term. P-value is represented by the color map.

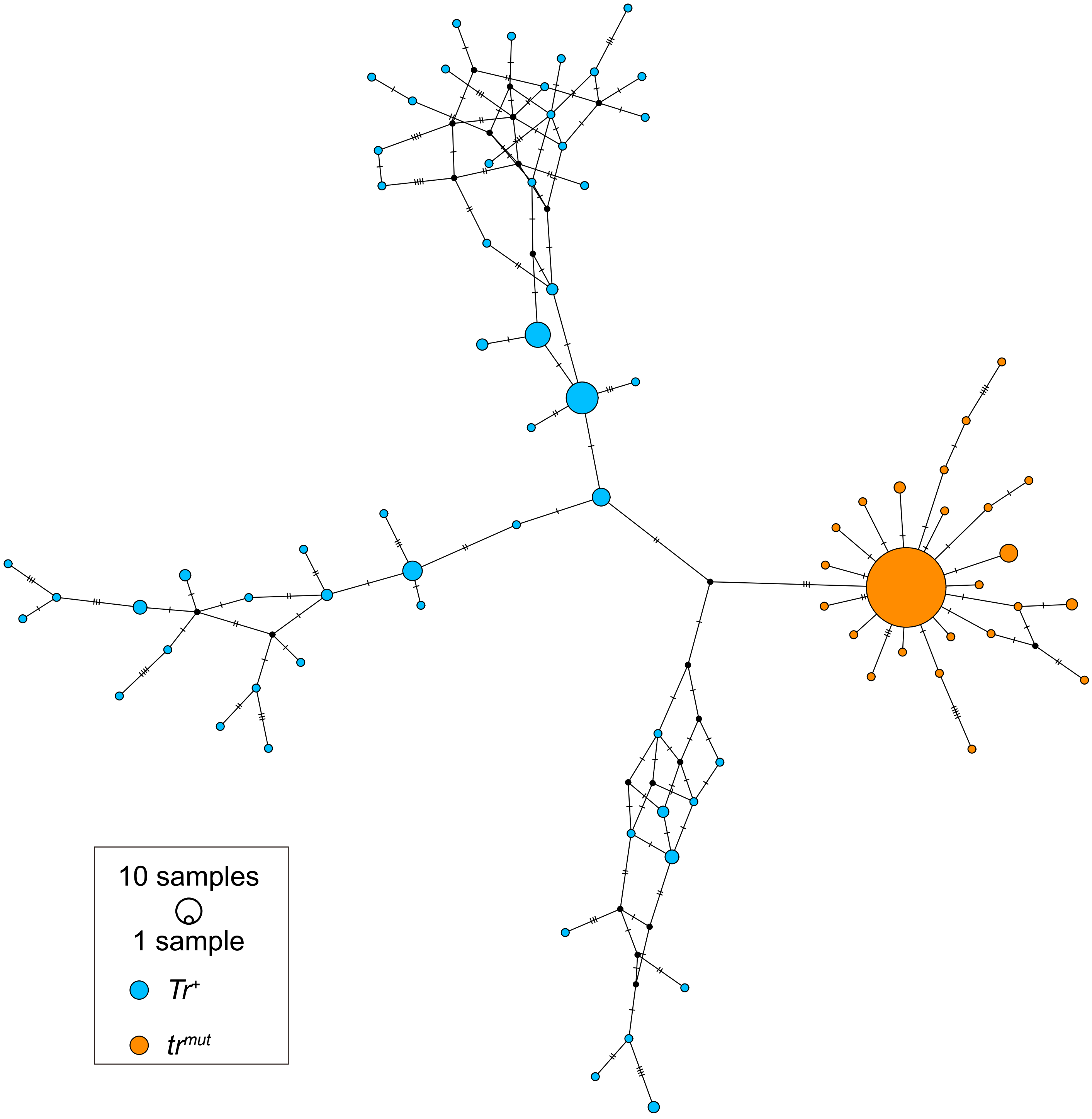


Figure S6. Haplotype network of 55 *Tr*$^{+}$ and 24 *tr*$^{mut}$ haplotypes generated from 9 Kb nonrecombining *Tr* region from 139 domestic pigeons (35 fancy pigeons, 2 feral pigeons, and 102 racing pigeons). The mutations was showed by hatch marks. The orange and blue circles represent *tr*$^{mut}$ and *Tr*$^{+}$ haplotypes, respectively.

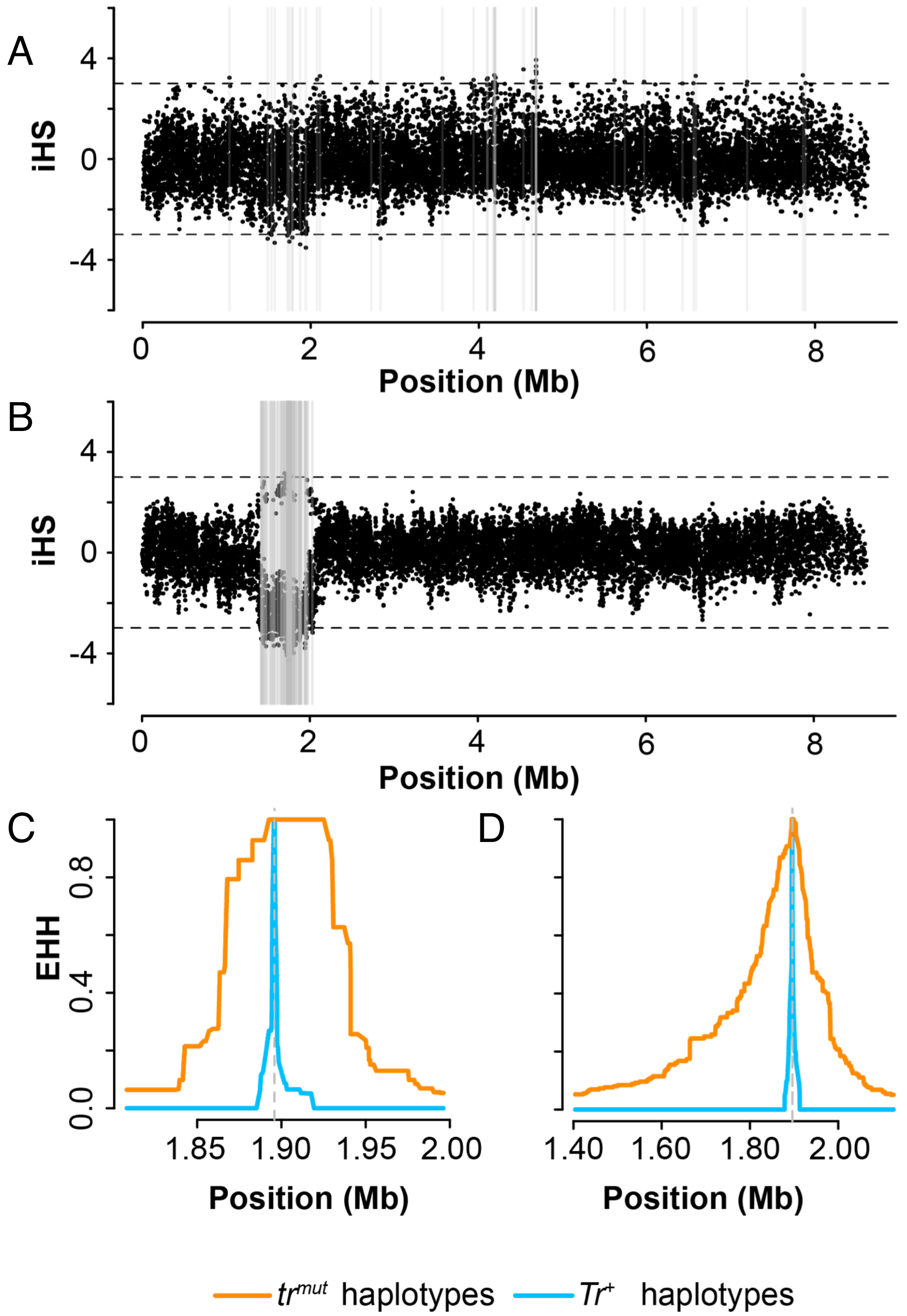


Figure S7. Selection analysis of haplotypes containing W49X mutation (*tr*$^{mut}$) and wild-type haplotypes (*Tr*$^{+}$) surrounding the *Tr* locus. Integrated haplotype score (iHS) was calculated for scaffold AKCR02000030.1 in fancy pigeons (A) and racing pigeons (B). The gray lines represent significance of absolute iHS scores of 3 or greater. Extended haplotype homozygosity (EHH) decay across the *Tr* locus region in fancy pigeons (C) and racing pigeons (D).

| | | | S32A | I33V | | | A36S | | | | | | | R52C | Y53F | | | | L57I | | | | | L98M | | | | L202F | | | | W261C | | | | V312A | | | | R337K | |
|---|---|---|---|---|---|---|---|---|---|---|---|---|---|---|---|---|---|---|---|---|---|---|---|---|---|---|---|---|---|---|---|---|---|---|---|---|---|---|---|---|---|
| | 30 | 31 | 32 | 33 | 34 | 35 | 36 | 37 | 38 | … | 49 | 50 | 51 | 52 | 53 | 54 | 55 | 56 | 57 | 58 | 59 | … | 97 | 98 | 99 | … | 201 | 202 | 203 | … | 260 | 261 | 262 | … | 311 | 312 | 313 | … | 336 | 337 | 338 |
| *Columba livia* | N | V | S | I | I | N | A | P | T | | W | N | S | R | Y | H | K | E | L | N | P | | L | L | V | | Q | L | L | | P | W | Q | | Y | V | T | | R | R | V |
| *Gallus gallus* | . | . | . | . | . | . | . | . | . | | . | A | . | . | . | . | V | . | . | . | . | | . | . | M | | . | . | . | | . | . | . | | . | . | . | | . | . | Y |
| *Amazona vittata* | . | . | . | . | . | . | . | . | . | | . | T | N | C | . | . | . | A | . | . | . | | . | . | M | | . | . | . | | . | . | . | | . | . | . | | . | . | Y |
| *Buceros rhinoceros* | . | . | . | . | . | . | . | . | . | | . | T | . | . | . | . | . | . | I | . | . | | . | . | M | | . | . | . | | . | . | . | | . | . | . | | . | . | Y |
| *Charadrius vociferus* | . | . | . | . | . | . | . | . | . | | . | T | . | . | . | . | . | . | . | . | . | | . | . | M | | . | . | . | | . | . | . | | . | . | . | | . | . | C |
| *Cuculus canorus* | . | . | . | . | . | . | S | . | . | | . | T | . | . | . | . | . | D | . | S | . | | . | . | M | | . | . | V | | . | . | . | | . | . | . | | . | . | Y |
| *Egretta garzetta* | . | . | . | . | . | . | . | . | . | | . | A | . | . | F | . | . | . | . | . | . | | . | . | M | | . | . | . | | . | . | . | | . | A | . | | . | . | C |
| *Leptosomus discolor* | . | . | . | . | . | . | . | . | . | | . | S | . | . | . | . | . | . | . | . | . | | . | . | M | | . | . | V | | . | . | . | | . | . | . | | . | . | Y |
| *Nestor notabilis* | . | . | . | . | . | . | . | . | . | | . | T | N | . | . | . | . | A | . | . | . | | . | . | M | | . | . | . | | . | C | H | | . | . | . | | . | . | Y |
| *Parotia lawesii* | . | . | . | V | . | . | . | . | . | | . | S | . | . | . | . | . | . | . | . | . | | . | . | M | | . | F | . | | . | . | . | | . | . | . | | . | . | C |
| *Phaethon lepturus* | . | . | . | . | . | . | . | . | . | | . | T | . | . | . | . | . | . | . | S | L | | . | . | M | | . | . | . | | . | . | . | | . | . | . | | . | K | Y |
| *Phoenicopterus ruber* | . | . | . | . | . | . | . | . | . | | . | T | . | . | . | . | . | . | . | . | . | | . | . | M | | . | . | . | | . | . | . | | . | . | . | | . | . | Y |
| *Pterocles gutturalis* | . | . | A | . | . | . | . | . | . | | . | T | . | . | . | R | . | . | . | . | . | | . | . | M | | . | . | V | | . | . | . | | . | . | . | | . | . | C |
| *Streptopelia turtur* | . | . | . | . | . | . | . | . | . | | . | S | I | . | . | . | . | . | . | . | . | | . | . | . | | . | . | . | | . | . | . | | . | . | . | | . | . | . |
| *Struthio camelus* | . | . | . | . | . | . | . | . | . | | . | T | . | . | . | . | . | K | . | . | . | | . | M | M | | . | . | I | | . | . | . | | . | . | . | | . | . | H |
| *Phalacrocorax carbo* | . | . | . | . | . | . | . | . | . | | . | T | . | . | . | . | . | . | . | . | . | | . | . | M | | . | . | . | | . | . | . | | . | . | . | | . | . | S |
| *Nannopterum auritus* | . | . | . | . | . | . | . | . | . | | . | T | . | . | . | . | . | . | . | . | . | | . | . | M | | . | . | . | | . | . | . | | . | . | . | | . | . | S |
| *Nannopterum brasilianus* | . | . | . | . | . | . | . | . | . | | . | T | . | . | . | . | . | . | . | . | . | | . | . | M | | . | . | . | | . | . | . | | . | . | . | | . | . | S |
| *Nannopterum harrisi* | . | . | . | . | . | . | . | . | . | | . | T | . | . | . | . | . | . | . | . | . | | . | . | M | | . | . | . | | . | . | . | | . | . | . | | . | . | S |
| *Urile pelagicus* | . | . | . | . | . | . | . | . | . | | . | T | . | . | . | . | . | . | . | . | . | | . | . | M | | . | . | . | | . | . | . | | . | . | . | | . | . | S |
| *Alligator mississippiensis* | . | . | . | . | . | . | . | . | . | | . | R | . | . | . | Q | M | . | . | . | . | | . | . | . | | . | . | . | | . | . | . | | . | . | . | | . | . | F |
| *Pelodiscus sinensis* | . | . | . | . | . | . | . | . | . | | . | S | . | . | . | Q | T | . | . | . | T | | . | . | . | | . | . | . | | . | . | . | | . | . | . | | . | . | F |
| *Salvator merianae* | . | . | . | . | . | S | . | . | . | | . | W | N | . | . | Q | A | . | . | S | . | | . | . | M | | . | . | . | | . | . | H | | . | . | . | | . | . | . |

| | | S357C | | | | | | | Y364H | | | | | | | | | | | | F376L | | | | I409V | | | | | S414N | | | T484Tfs3 | | | | | | | | |
|---|---|---|---|---|---|---|---|---|---|---|---|---|---|---|---|---|---|---|---|---|---|---|---|---|---|---|---|---|---|---|---|---|---|---|---|---|---|---|---|---|---|
| | 356 | 357 | 358 | 359 | 360 | 361 | 362 | 363 | 364 | 365 | 366 | 367 | 368 | 369 | 370 | 371 | 372 | 373 | 374 | 375 | 376 | 377 | … | 408 | 409 | 410 | 411 | 412 | 413 | 414 | 415 | … | 484 | 485 | 486 | 487 | 488 | 489 | 490 | 491 | 492 |
| *Columba livia* | L | S | L | T | Y | Q | E | L | Y | S | W | V | S | Y | V | S | M | T | S | I | F | A | | M | I | G | G | T | V | S | W | | T | E | L | Y | E | R | R | E | L |
| *Gallus gallus* | F | . | . | . | . | . | . | . | . | . | . | . | P | . | . | . | . | M | . | . | . | . | | . | . | . | . | . | . | . | . | | . | . | . | N | . | . | S | Q | . |
| *Amazona vittata* | F | . | . | . | . | . | . | . | . | P | . | . | P | . | . | . | . | . | . | . | . | . | | . | . | . | . | . | I | . | . | | . | . | . | . | . | . | . | Q | . |
| *Buceros rhinoceros* | F | . | . | . | . | . | . | . | . | . | . | . | P | . | . | . | . | . | . | . | . | . | | A | V | . | . | . | . | . | . | | . | . | . | . | . | . | . | Q | . |
| *Charadrius vociferus* | F | . | . | . | . | . | D | . | . | . | . | . | P | . | . | . | . | . | . | . | L | . | | . | . | A | . | . | I | . | . | | . | . | . | . | . | . | . | Q | . |
| *Cuculus canorus* | F | . | . | . | . | . | . | . | . | . | . | . | P | . | . | . | . | . | A | . | . | . | | . | . | A | . | . | I | . | . | | N | . | . | . | . | . | . | Q | . |
| *Egretta garzetta* | F | . | . | . | . | . | . | . | . | . | . | . | P | . | . | . | . | V | . | L | . | . | | . | . | . | . | . | I | . | . | | . | . | . | . | . | . | . | Q | . |
| *Leptosomus discolor* | F | C | . | . | . | . | . | . | . | L | . | . | P | . | . | . | . | A | . | . | . | . | | . | . | . | . | . | I | . | . | | . | . | . | . | . | . | . | Q | . |
| *Nestor notabilis* | F | . | . | . | . | . | . | . | . | P | . | . | P | . | . | . | . | . | . | . | . | . | | . | . | . | . | . | I | . | . | | . | . | . | F | . | . | . | Q | . |
| *Parotia lawesii* | F | . | . | . | . | . | . | . | . | . | . | . | P | . | M | . | . | . | C | V | . | . | | . | . | A | . | . | . | . | . | | . | . | . | H | . | . | . | Q | . |
| *Phaethon lepturus* | F | . | . | . | . | . | . | . | . | . | . | . | P | . | . | . | . | . | . | . | . | . | | . | . | . | . | . | I | . | . | | . | . | . | . | . | . | . | Q | . |
| *Phoenicopterus ruber* | F | . | . | . | . | . | . | . | H | . | . | . | P | . | . | . | . | . | . | . | . | . | | . | . | . | . | . | I | . | . | | . | . | . | . | . | . | . | Q | . |
| *Pterocles gutturalis* | F | . | . | . | . | . | . | . | . | . | . | . | P | . | . | . | . | M | C | . | . | . | | A | . | . | . | . | I | . | . | | . | . | . | . | . | . | . | Q | . |
| *Streptopelia turtur* | F | . | . | . | . | . | . | . | . | . | . | . | P | . | . | . | . | . | . | . | . | . | | . | . | . | . | . | . | N | . | | . | . | . | . | . | . | . | Q | V |
| *Struthio camelus* | F | . | . | . | . | . | . | . | . | . | . | . | P | . | . | . | . | . | . | . | . | . | | . | . | . | . | A | I | . | . | | . | . | . | H | . | . | . | Q | . |
| *Phalacrocorax carbo* | F | . | . | . | . | . | . | . | . | . | . | . | P | . | . | . | V | . | . | L | . | . | | . | . | . | . | . | I | . | . | | . | A | . | * | - | - | - | - | - |
| *Nannopterum auritus* | F | . | . | . | . | . | . | . | . | . | . | . | P | . | . | . | V | . | . | L | . | . | | . | . | . | . | . | I | . | . | | . | A | . | * | - | - | - | - | - |
| *Nannopterum brasilianus* | F | . | . | . | . | . | . | . | . | . | . | M | P | . | . | . | V | . | . | L | . | . | | . | . | . | . | . | I | . | . | | . | A | . | * | - | - | - | - | - |
| *Nannopterum harrisi* | F | . | . | . | . | . | . | . | . | . | . | . | P | . | . | . | V | . | . | L | . | . | | . | . | . | . | . | I | . | . | | . | A | . | * | - | - | - | - | - |
| *Urile pelagicus* | F | . | . | . | . | . | . | . | . | . | . | . | P | . | . | . | V | . | . | L | . | . | | . | . | . | . | . | I | . | . | | . | A | . | * | - | - | - | - | - |
| *Alligator mississippiensis* | F | . | . | . | . | . | . | Q | . | . | . | . | P | . | L | . | L | A | . | . | . | . | | . | . | . | . | S | I | . | . | | P | . | . | C | D | . | I | Q | . |
| *Pelodiscus sinensis* | F | . | . | . | . | . | . | . | . | P | . | . | P | . | L | . | . | A | . | . | . | . | | . | . | . | . | S | . | . | . | | A | . | . | . | . | . | . | Q | . |
| *Salvator merianae* | F | . | . | . | . | . | - | V | . | . | . | A | P | . | L | . | . | A | C | . | . | . | | T | . | A | . | . | I | . | . | | R | . | . | C | . | T | Q | Q | . |

Figure S8. The missense and frame-shifting mutations of *SLC2A11B* in avian species. A total of 87 coding sequences of 70 Aves, 2 Crocodilia, 6 Testudines, and 9 Squamata were aligned. Partial alignment of amino acid sequences of SLC2A11B among avian with conserved species-specific missense mutations was shown. The species-specific missense mutations and frame-shifting mutation in cormorants are labeled at the top of table, and the deleterious mutations were marked in red.
